# IL-4-induced hysteresis in naïve T cell activation

**DOI:** 10.1101/2020.08.31.275842

**Authors:** Alexandre P. Meli, Yaqiu Wang, Dimitri A. de Kouchkovsky, Yong Kong, Malay K. Basu, Sourav Ghosh, Carla V. Rothlin

## Abstract

Naïve T cells are generally considered to be a homogeneous population, but for their unique T cell receptors (TCRs). Naïve T cells are activated within a specific cytokine milieu upon interaction with antigen-presenting cells through cognate TCR::MHC-peptide interaction and co-stimulation. Here we demonstrate that naïve T cells are transcriptionally heterogeneous, and that the relative proportions of transcriptionally distinct naïve T cell subpopulations are modified by immune responses, such as during helminth infection. Not only are cognate naïve T cells activated during an immune response, but the cytokine produced - such as IL-4 during helminth infection - changes the transcriptome of bystander naïve T cells. Such changes in gene expression and population level heterogeneity in bystander naïve T cells result in altered responses to a concurrent immune challenge, for instance, hypo-responsiveness to vaccination. Thus, naïve T cell activation is not the result of a singular temporal event, but is characterized by hysteresis. Our studies suggest that antigen-agnostic, cytokine-dependent naïve T cell conditioning and resulting hysteresis is a mechanism that integrates input signals from concurrent infections for the regulation of the overall magnitude of the immune response.

## Main text

TCR diversity generates unique naïve T cells that play an essential role in host defense against a large and unpredictable range of pathogens or transformed self. The conversion of a naïve T cell to an activated T cell commonly occurs in secondary lymphoid organs and requires two essential membrane-bound signals: i) engagement of the T cell receptor (TCR) by a specific antigenic peptide presented on the major histocompatibility complex (MHC) of the antigen-presenting cell (APC), and ii) interaction of the non-antigen-specific CD28 co-stimulatory cell surface molecule on the T cell with B7 ligands on the APC. Besides TCR sequence diversity, naïve T cells have largely been thought to represent a fairly homogeneous population in a quiescent state prior to antigen encounter. Recently, it has been recognized that naïve T cells are also characterized by transcriptional heterogeneity at steady state (*1*). Whether naïve T cell transcriptional heterogeneity is fixed or subjected to perturbation, and whether this diversity contributes to differential T cell activation, remains unknown.

In addition to the two membrane-bound signals, a third critical constituent of T cell activation is cytokines produced by the APCs or other cells. While TCR and co-stimulatory signals are restricted to the cognate APC::T cell interaction during a clonal priming event, cytokines are diffusible. Therefore, unlike the specificity of the membrane-restricted signals for the target T cell, bystander naïve T cells can also be the substrate of the diffusible signal (*2*). We hypothesized that bystander naïve T cells are conditioned by the cytokine milieu produced during an immune response. We further posited that this cytokine-mediated conditioning during an ongoing immune response alters the heterogeneity of bystander naïve T cells, and their subsequent activation, when encountering their cognate antigen.

### Helminth infection alters the transcriptional heterogeneity of naïve CD4^+^ T cells

To test our hypothesis, we infected mice with *Nippostongylus brasiliensis* (*N. brasiliensis*) wherein a type 2-biased immune response is elicited in the draining lymph nodes characterized by production of cytokines IL-4, IL-5 and IL-13 (*3*). We sorted CD3^+^CD4^+^CD62L^+^CD44^−^B220^−^CD25^−^CXCR5^−^PD1^−^IL-4-GFP^−^IL-21-Kat^−^ cells (naïve CD4^+^ T cells) from uninfected mice and from mice 1 week after they were infected with *N. brasiliensis*, when IL-4 levels are high (*3, 4*). Additionally, we sorted T_H_2 cells from *N. brasiliensis*-infected mice 1 week after helminth infection. We performed single-cell RNA sequencing (scRNAseq) on these populations. As expected, we observed a population of T_H_2 cells characterized by *Gata3* expression in *N. brasiliensis*-infected mice (**Fig. S1A, C**). Naïve CD4^+^ T cells from uninfected mice clustered primarily into four distinct subsets (**Fig. 1A**). Jaccard similarity co-efficient analyses indicated that our subset 4 approximated the previously characterized TCR activation and pathway cluster (*1*) (**Fig. S1B**). Similarly, our subset 3 approximated the previously characterized type I IFN signature-enriched cluster (*1*) (**Fig. S1B**). Importantly, we observed clear differences in the population level heterogeneity in naïve CD4^+^ T cells isolated from uninfected *versus N. brasiliensis*-infected mice (**Fig. 1A-B**). A unique population of naïve CD4^+^ T cells (cluster 2) emerged in mice following *N. brasiliensis* infection (**Fig. 1A-B**). This heterogeneity was independent of T_H_2 markers (**Fig. S1C**). Cell lineage and pseudotime inference using Slingshot (*5*) indicated that cluster 1 and cluster 0 are the primordial clusters, while clusters 2, 3 and 4 arise later in time from cluster 1 *via* cluster 0 (**Fig. 1C**). Trajectory analysis further indicated that clusters 2, 3 and 4 arise independently of each other (**Fig. 1D**). Consistent with this observation, diffusion map (*6*) reconstruction also revealed three distinct trajectories for clusters 2, 3 and 4 (**Fig. S1D**). The 20 most highly upregulated genes in cluster 2, relative to all other clusters, associated with an IL-4/IL-13 signaling pathway as the top category using the Reactome Pathway Knowledgebase (*7*) (*p* value = 1.26×10^−5^) (**Fig. 1E**). Of these 20 upregulated genes, 14 of the genes, including *Il4ra*, have been previously reported to be induced in T cells by IL-4 (*8*), a prototypical cytokine produced during helminth infection. By contrast, the top 20 downregulated genes in cluster 2 relative to most other clusters associated with the IFNα/β-signaling category in the Reactome Pathway Knowledgebase (*p* value = 3.04×10^−9^) (**Fig. 1E**). Interestingly, IL-4 has been shown previously to suppress type I IFN inducible genes in dendritic cells and T cells (*8, 9*), suggesting that the dynamic changes in naïve T cell heterogeneity were driven by IL-4.

**Figure 1.**
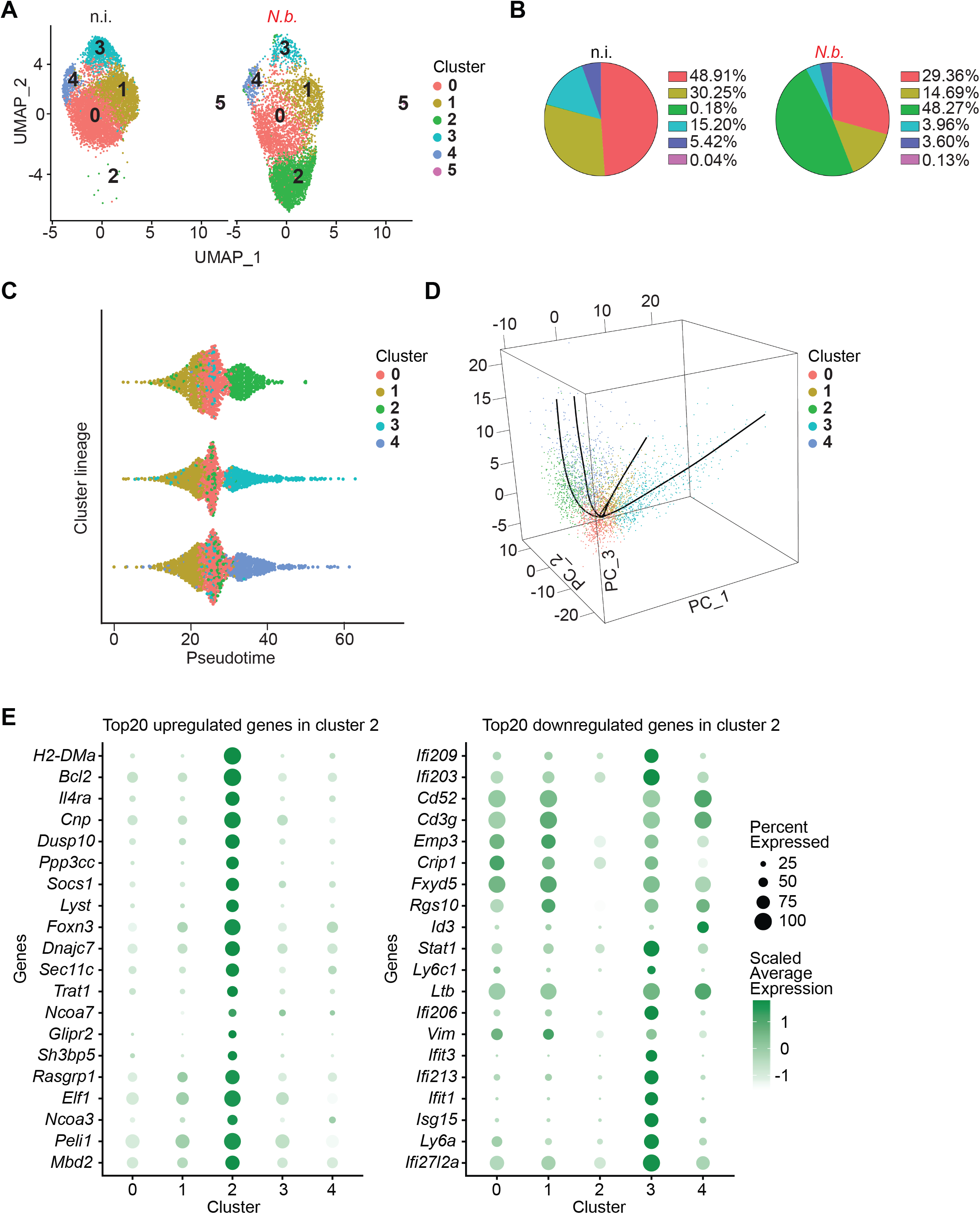
Increased transcriptional heterogeneity of naïve CD4^+^ T cells upon helminth infection. scRNA sequencing of naïve CD3^+^CD4^+^CD62L^+^CD44^−^B220^−^CD25^−^CXCR5^−^PD1^−^IL-4-GFP^−^IL-21-Kat^−^ T cells FACS-sorted from mesenteric lymph nodes (LNs) of *N. brasiliensis (N.b.)*-infected (day 7) and non-infected (n.i.) mice. (**A**) Uniform Manifold Approximation and Projection (UMAP) of single naïve T cells. Each dot represents an individual cell and is colored according to cell cluster. (**B**) Pie chart depicting the percentage of naïve T cells assigned to each cluster. (**C**) Cluster lineages identified by Slingshot. The x-axis represents pseudotime and the y-axis represents the 3 lineages identified by Slingshot. Each dot represents a cell, and dots are colored as in A. Note, cluster “5”, which contains only a few cells, has been removed from the analysis. (**D**) 3D scatter plot showing the 3 trajectories identified in C. x, y, and z axes represent principal components 1, 2, 3, respectively. Dots are cells and colored as in C. The common root was identified in cluster 1, which gives rise to cluster 0. All other clusters are found to arise from cluster 0. (**E**) Top20 upregulated and Top20 downregulated genes in cluster 2 in comparison to other indicated clusters. Representative of 1 experiment, with single cells pooled from 4 mice per group.

### Helminth-induced changes in naïve CD4^+^ T cells are driven by IL-4, and correlate with reduced amplitude of the immune response to vaccination

Analysis of scRNAseq data demonstrated that IL-4Rα and γ_c_ (common γ chain), but not IL-5Rα or IL-13Rα1, were expressed in naïve CD4^+^ T cells (**Fig. S2A**). Therefore, we hypothesized that IL-4, but not the other prototypical helminth infection-induced cytokines IL-5 or IL-13, pre-primes bystander naïve T cells during helminth infection. To test our hypothesis, we first performed RNAseq analysis of sorted naïve CD4^+^ T cells from uninfected and *N. brasiliensis*-infected mice, and also from uninfected mice injected with IL-4c. 98 genes were differentially upregulated at least 2-fold in naïve CD4^+^ T cells isolated from *N. brasiliensis*-infected mice *versus* naïve CD4^+^ T cells isolated from uninfected mice (**Fig. 2A**). We validated the induction of selected IL-4-dependent genes including *Pros1*, *Lgals3*, *Ppp3cc* and *Aqp9* by qRT-PCR (**Fig. S2B**). Furthermore, naïve CD4^+^ T cells isolated from uninfected mice treated with IL-4c displayed upregulation of 242 genes in comparison to uninfected, untreated mice (**Fig. S3**). Of the *N. brasiliensis*-upregulated genes, 85.7% (84 genes) were shared with naïve CD4^+^ T cells isolated from IL-4c-treated mice (**Fig. 2B**). Only 14.3% genes (14 genes) were uniquely upregulated in naïve CD4^+^ T cells isolated from *N. brasiliensis*-infected mice (**Fig. 2B**). Hypergeometric test revealed that the overlap of upregulated genes between *N. brasiliensis*-infected or IL-4c-treated mice was statistically significant. This overlap suggests that naïve CD4^+^ T cells are transcriptionally reprogrammed by IL-4 during helminth infection.

**Figure 2.**
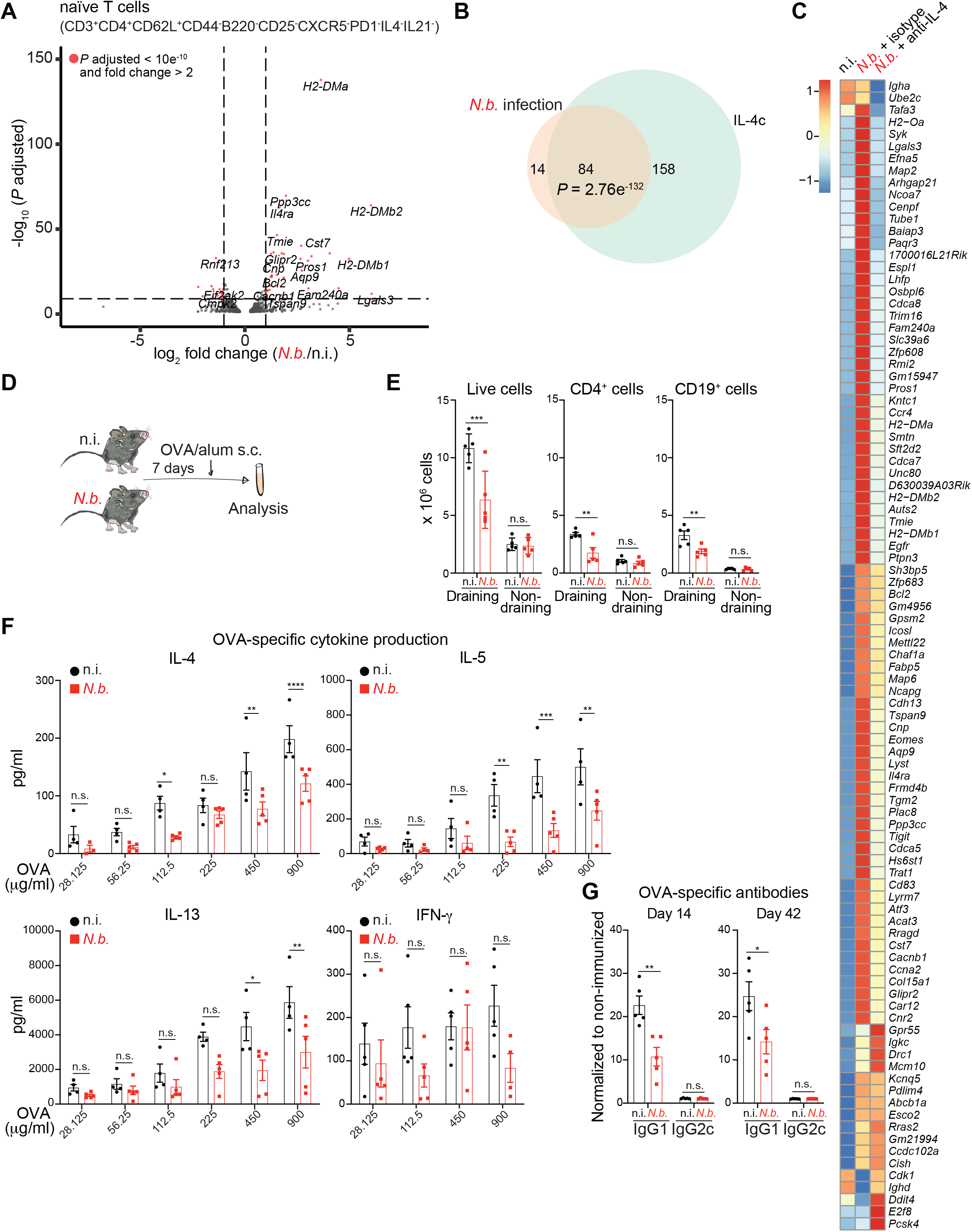
Transcriptional re-programming of naïve T cells is induced by IL-4 and correlates with reduced response to vaccines in helminth-infected mice. (**A**) Naïve (CD3^+^CD4^+^CD62L^+^CD44^−^B220^−^CD25^−^ CXCR5^−^PD1^−^IL-4-GFP^−^IL-21-Kat^−^) T cells were sorted from mesenteric lymph nodes (LNs) from *N. brasiliensis* (*N.b.*)-infected (day 7) or non-infected (n.i.) mice. Volcano plot represents genes differentially regulated at least 2-fold in *N.b*-infected mice compared to non-infected controls, as determined by RNAseq. (**B**) Naïve (CD3^+^CD4^+^CD62L^+^CD44^−^B220^−^CD25^−^CXCR5^−^PD1^−^IL-4-GFP^−^IL-21-Kat^−^) T cells were sorted from mesenteric LNs of *N.b.*-infected (day 7), 1 day after IL-4 complex (IL-4c) administration, or non-infected mice. Venn diagrams depict overlap in genes at least 2-fold upregulated in naïve CD4^+^ T cells from *N.b.*-infected or IL-4c-treated mice in comparison to naïve CD4^+^ T cells from non-infected mice. A-B are representative of 2 independent experiments each with samples pooled from 4 mice per group. Analyzed using hypergeometric test. (**C**) Heat map depicting the expression of the genes upregulated at least 2-fold at 1 week post-*N.b.* infection shown in A and B. Naïve CD3^+^CD4^+^B220^−^CD25^−^CD44^−^CD62L^+^ T cells were sorted from mesenteric LNs of n.i. mice and *N.b.*-infected (day 7) mice treated with IL-4 neutralizing antibody or isotype control every 48 hours beginning 2 days before infection. Gene expression was determined by RNAseq. Representative of 1 experiment with pooled samples from 4 mice per group. (**D**) Experimental scheme for E to G. n.i. controls or mice infected with *N.b.* for 7 days were immunized with ovalbumin(OVA)/alum subcutaneously in the footpad. (**E**) Number of total, CD4^+^CD19^−^ and CD4^−^CD19^+^ cells from draining and non-draining LNs collected 7 days post-immunization. (**F**) Concentrations of indicated cytokines in the supernatant of enriched CD4^+^ T cells from draining LNs of immunized animals that were re-stimulated for 96 hours with OVA in the presence of irradiated splenocytes. (**G**) OVA-specific antibody concentrations for the indicated isotypes in the sera of immunized mice collected at the indicated times post-immunization. E to G are representative of 3 independent experiments and each data point represents a unique animal. Data in E and G were analyzed using two-tailed Student’s t-test. Data in F were analyzed using 2-way ANOVA with Holm-Sidak’s multiple comparisons test. ****p<0.0001 ***p<0.001, **p<0.01 and *p<0.05. n.s. is non-significant. Error bars, SEM.

Next, we performed RNAseq on naïve CD4^+^ T cells isolated from *N. brasiliensis*-infected mice and from *N. brasiliensis*-infected mice injected with an IL-4 neutralizing antibody. IL-4 was effectively neutralized by antibody treatment (**Fig. S4A**). Of the 98 genes induced by *N. brasiliensis* infection in Fig. 2A and B, 78 (79.6%) genes were downregulated by IL-4 blockade (**Fig. 2C**). This downregulated set included well known IL-4-inducible genes such as *Pros1*, *Il4ra* and *Ppp3cc.* Furthermore, we validated the downregulation of *Pros1*, *Lgals3*, *Tgm2* and *Aqp9* upon neutralization of IL-4 by qRT-PCR (**Fig. S4B**). These results conclusively demonstrate that the bulk of the transcriptional upregulation in naïve CD4^+^ T cells following *N. brasiliensis* infection is driven by IL-4.

To examine the functional consequences of this alteration in the naïve T cell transcriptome, if any, we performed foot-pad immunization with ovalbumin (OVA) and the adjuvant alum that favors a type 2 immune response, in uninfected mice and in *N. brasiliensis*-infected mice 1 week post-infection (**Fig. 2D**). We observed a significantly lower total cellularity of the foot-pad draining lymph nodes as well as number of CD4^+^ T and B cells in vaccinated, *N. brasiliensis*-infected mice in comparison to vaccinated, uninfected mice (**Fig. 2E**). To confirm if the quality of antigen-specific CD4^+^ T cell responses in vaccinated *N. brasiliensis*-infected mice was also impaired, total CD4^+^ T cells were enriched from popliteal and inguinal draining lymph nodes and re-stimulated *in vitro* in the presence of irradiated splenocytes with increasing concentrations of OVA. CD4^+^ T cells isolated from vaccinated, *N. brasiliensis*-infected hosts exhibited decreased antigen sensitivity in producing the expected T_H_2-associated cytokines IL-4, IL-5 and IL-13 (**Fig. 2F**). T-dependent antibody responses to OVA-alum immunization were also impaired after *N. brasiliensis* infection as exhibited by a decrease in OVA-specific IgG1 (**Fig. 2G**). Of note, we did not observe a skewing towards a type 1 response since neither the production of IFN-γ by CD4^+^ T cells nor the amount of IgG2c in the serum was increased in vaccinated, *N. brasiliensis*-infected mice relative to vaccinated, uninfected mice. (**Fig. 2F-G**). Moreover, we extended our investigation of the suppression of the magnitude of the immune response following helminth infection to an independent model where mice were infected with *Heligmosomoides polygyrus bakeri* (*H. polygyrus bakeri*). Infection with this helminth similarly attenuated the OVA-specific cellular and humoral responses, recapitulating our observations with *N. brasiliensis* infection (**Fig. S5**). Furthermore, the IL-4-driven gene *Pros1* was upregulated in naïve CD4^+^ T cells isolated from *H. polygyrus bakeri*-infected mice (**Fig. S6**). These results suggest that helminth infection and IL-4-induced reprogramming of naïve CD4^+^ T cells and changes in population heterogeneity resulted in a reduced immune response following concurrent challenges.

To test if helminth infection uniquely impairs the OVA-alum driven type 2 responses, or if this effect can be extended beyond this specific effector subtype, we immunized *N. brasiliensis*-infected mice with OVA-CpG (**Fig. S7A**), which results in a mixed type 1 and 17 effector response characterized by the differentiation of IFN-γ- and IL-17-producing CD4^+^ T cells, respectively. Not only did prior infection with *N. brasiliensis* result in a lower total number of cells in the footpad-draining lymph nodes 1 week after vaccination in comparison to vaccinated, non-infected mice, but also the number of CD4^+^ T cells and B cells was decreased (**Fig. S7B**). The sensitivity of the antigen-specific response of IFN-γ- and IL-17-producing CD4^+^ T cells was also significantly impaired by *N. brasiliensis* infection (**Fig. S7C**). A significantly reduced concentration of OVA-specific IgG2c was detected in vaccinated, *N. brasiliensis*-infected mice in comparison to vaccinated, non-infected mice, demonstrating that antigen-specific antibody responses were also diminished in *N. brasiliensis*-infected mice (**Fig. S7D**). Taken together, our results indicate that helminth infections induce IL-4-dependent transcriptional changes in bystander naïve T cells, and these transcriptional changes correlate with a diminished magnitude of subsequent antigen-specific immune responses.

### Helminth infection pre-primes a naïve CD4^+^ T cell hypo-responsive state

To test if the changes in naïve CD4^+^ T cells can causally account for the observed changes in the magnitude of the immune response, we examined if naïve CD4^+^ T cells from either uninfected or *N. brasiliensis*-infected mice had equivalent capacity for proliferation, or if naïve T cells from *N. brasiliensis*-infected mice were conditioned to be hypo-responsive. To this end, naïve T cells were enriched from uninfected OT-II TCR transgenic mice or from OT-II TCR transgenic mice 7 days post-infection with *N. brasiliensis*. These cells were labelled with CellTrace^TM^ Violet dye and transferred into uninfected or *N. brasiliensis*-infected congenic CD45.1 recipient mice, respectively, 24 hours prior to immunization with OVA-alum (**Fig. 3A**). By 4 days post immunization, proliferating OT-II CD4^+^ T cells were found in the footpad-draining lymph nodes in both uninfected and *N. brasiliensis*-infected mice (**Fig. 3B-C**). CD4^+^ T cells transferred from *N. brasiliensis*-infected donors to *N. brasiliensis*-infected hosts proliferated markedly less compared to CD4^+^ T cells isolated and transferred into uninfected counterparts. Therefore, the immune environment in helminth-infected hosts impaired CD4^+^ T cell proliferation (**Fig. 3B-C**). Of note, helminth infection did not alter the number of OVA^+^ activated dendritic cells in the draining lymph nodes (**Fig. S8A-C**) or T-cell-independent antibody responses (**Fig. S8D**), suggesting that the diminished immune response to vaccination is not due to impaired dendritic cell or B cell function.

**Figure 3.**
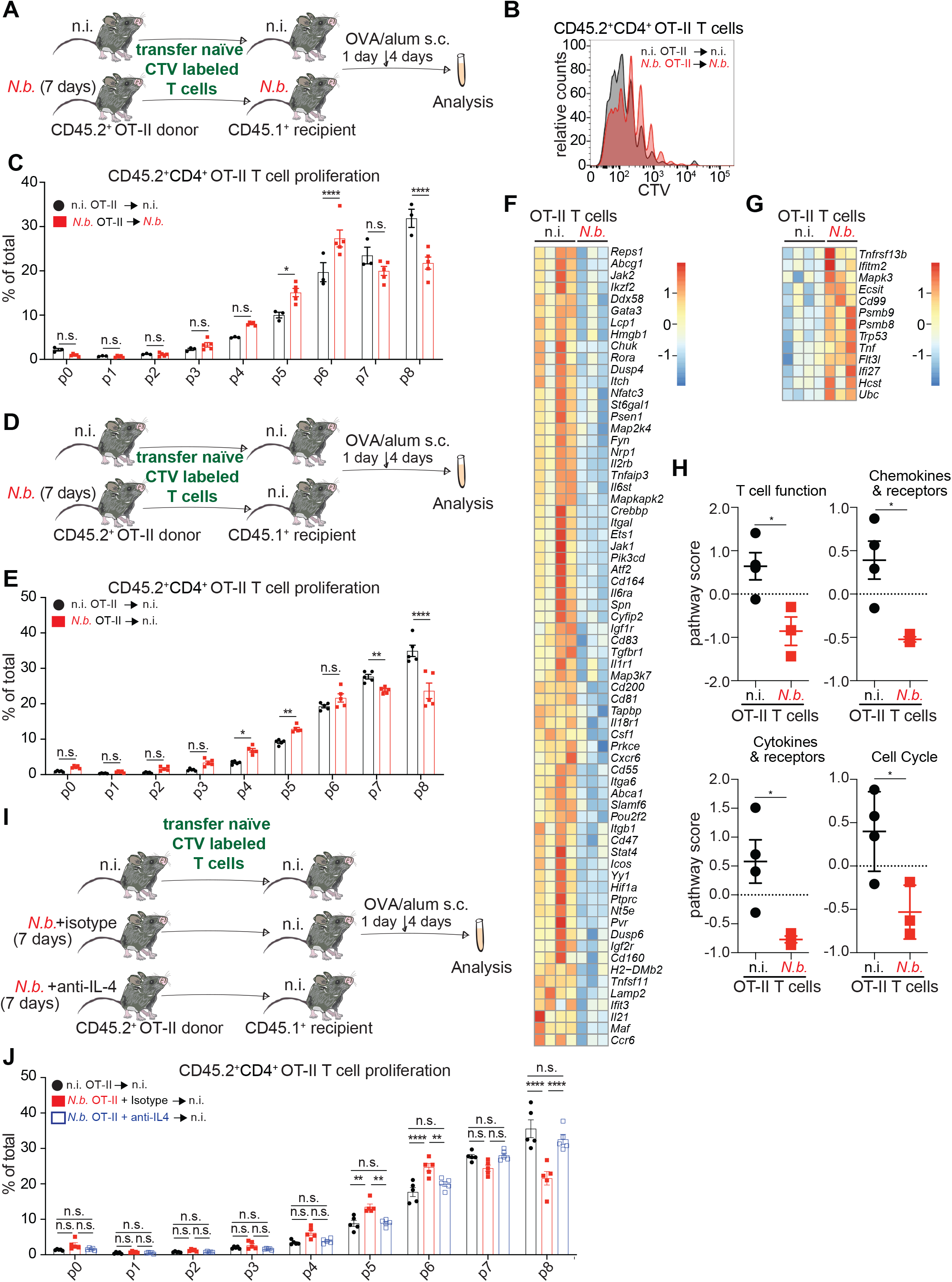
IL-4 pre-primes naïve CD4^+^ T cell to a hypo-responsive state. (**A**) Experimental scheme for B and C. Naïve CD4^+^ T cells were enriched from spleens of non-infected (n.i.) or 7 day *N. brasiliensis* (*N.b.*)- infected CD45.2^+^ OT-II transgenic mice, labeled with cell trace violet (CTV) and transferred into n.i. or 7 day *N.b.*-infected CD45.1 wild type recipients. Mice were immunized 24 hours after OT-II T cell transfer with OVA-alum and proliferation of OT-II T cells isolated from draining lymph nodes (LNs) was assessed 4 days later. (**B**) Representative histograms of CTV dilution of CD45.2^+^CD4^+^ T cells. (**C**) Quantification of OT-II T cell proliferation. (**D**) Experimental scheme for E-H. Naïve CD4^+^ T cells were enriched from spleens of n.i. and 7 day *N.b.*-infected CD45.2^+^ OT-II transgenic mice, labeled with CTV (for proliferation analysis shown in E) and transferred into n.i. CD45.1^+^ recipients. Mice were immunized 24 hours after OT-II T cell transfer with OVA/alum and analysis were performed 4 days later on cells isolated from draining LNs. (**E**) Quantification of OT-II T cell proliferation. (**F** and **G**) Heat map depicting genes significantly downregulated (F) or upregulated (G) in CD45.2^+^CD4^+^ OT-II T cells FACS-sorted from draining LNs and quantified using the nCounter PanCancer Immune Profiling Panel and the nCounter system (NanoString Technologies). (**H**) Pathway score of the dataset represented in F and G. (**I**) Experimental scheme for J. Naïve CD4^+^ T cells were enriched from spleens of n.i. and 7 day *N.b.*-infected CD45.2^+^ OT-II transgenic mice treated with IL-4 neutralizing antibody or isotype control, labeled with CTV and transferred into n.i. CD45.1^+^ recipients. Mice were immunized 24 hours after OT-II T cell transfer with OVA-alum and proliferation of OT-II T cells isolated from draining LNs was assessed 4 days later. (**J**) Quantification of OT-II T cell proliferation. Data in B,C and E are representative of two independent experiments. F-H are representative of one experiment with samples pooled from 3 mice per replicate. Data in J are representative of two independent experiments. Data in C, E and J were analyzed using 2-way ANOVA with Holm-Sidak’s multiple comparisons test. Data in H were analyzed using two-tailed Student’s t-test. Each data point represents a unique animal. ****p<0.0001 ***p<0.001, **p<0.01 and *p<0.05. n.s. is non-significant. Error bars, SEM.

To investigate if helminth infection can intrinsically alter naïve CD4^+^ T cells to differentially respond when challenged by cognate antigen, OT-II CD4^+^ T cells were isolated from uninfected or *N. brasiliensis*- infected CD45.2 mice and transferred into uninfected CD45.1 recipient mice (**Fig. 3D**). Of note, an equivalent number of OT-II CD4^+^ T cells were found in the popliteal lymph nodes 24 h post transfer (**Fig. S8E**). Following immunization, OT-II CD4^+^ T cells derived from *N. brasiliensis*-infected mice proliferated to a lesser extent compared to OT-II CD4^+^ T cells collected from uninfected donors (**Fig. 3E**). Therefore, helminth infection can pre-prime naïve CD4^+^ T cells to a cell-intrinsic hypo-responsive state.

Next, we investigated the T cell intrinsic transcriptional changes in activated T cells that have been pre-primed during helminth infection. To this end, RNA was isolated from transferred T cells 4 days after immunization and a focused gene expression pattern was analyzed using the nCounter PanCancer Immune Profiling Panel and the nCounter NanoString platform (NanoString Technologies). 81 genes were differentially expressed (p<0.05) between the CD45.2 OT-II CD4^+^ T cells transferred from uninfected or *N. brasiliensis*- infected mice (**Fig. 3F-G**). Specifically, 68 genes were significantly reduced in their expression in T cells transferred from *N. brasiliensis*-infected mice in comparison to those transferred from uninfected mice (**Fig. 3F**). Examples include *Gata3*, *Il2rb*, *Icos*, *Il21* and *Maf.* 13 genes were expressed in higher amounts in T cells transferred from *N. brasiliensis*-infected mice in comparison to those transferred from uninfected mice (**Fig. 3G**). An example includes the cell cycle arrest gene *Trp53* (**Fig. 3G**). Overall, pathways associated with T cell function, chemokines and their receptors, cytokines and their receptors, and cell cycle were expressed at significantly lower amounts in T cells transferred from *N. brasiliensis*-infected mice in comparison to those transferred from uninfected mice (**Fig. 3H**).

As the majority of transcriptional changes in naïve CD4^+^ T cells during helminth infection were IL-4-dependent, we hypothesized that IL-4 pre-priming during *N. brasiliensis* infection impaired the ability of naïve CD4^+^ T cells to proliferate in response to vaccination. To validate this hypothesis, we performed similar OT-II transfer experiments as previously described, along with an additional arm wherein IL-4 neutralizing antibodies were administered to the infected donors (**Fig. 3I**). Consistent with results shown in Fig. 3E, transferred naïve CD4^+^ T cells from uninfected OT-II donors proliferated significantly more in uninfected recipients after vaccination compared to naïve CD4^+^ T cells isolated from isotype-treated *N. brasiliensis*-infected mice transferred into uninfected recipients (**Fig. 3J**). Interestingly, IL-4 blockade in *N. brasiliensis*-infected OT-II donors restored the proliferative capacity of naïve T cells when these were transferred into uninfected hosts and recipient mice were vaccinated (**Fig. 3J**). These results demonstrate that helminth infection pre-primes naïve CD4^+^ T cells to a hypo-responsive state in an IL-4-dependent manner. Therefore, their activation is reduced when subsequently challenged by vaccination. Hysteresis is a phenomenon wherein the output of a system depends not only on its current inputs, but also on past inputs. Thus, naïve CD4^+^ T cell activation is subject to hysteresis wherein the passive exposure of bystander naïve CD4^+^ T cells to cytokines during helminth infection alters their response threshold to their cognate antigen.

### Hysteresis in naïve CD4^+^ T cell activation is rate-dependent

Hysteresis can be either rate-dependent, wherein the output decays as the input is reduced to zero, or rate-independent, which is an irreversible change in the state of the system after pre-priming. We performed scRNAseq to determine if the induced heterogeneity of naïve CD4^+^ T cells required tonic exposure to the helminth-elicited environment or if it was sustained after clearance of the helminth. Naïve CD4^+^ T cells were sorted from uninfected mice and from mice 1 week and 3 weeks after *N. brasiliensis* infection. The 3 week timepoint corresponds to complete elimination of the helminth. Naïve CD4^+^ T cells isolated from uninfected mice and from mice 1 week after *N. brasiliensis* infection recapitulated the appearance of cluster 2 after infection (**Fig. 4A-B**). By 3 weeks post-infection, cluster 2 dramatically contracted (**Fig. 4A-B**). By contrast, cluster 3, which was reduced at 1 week after *N. brasiliensis* infection, was restored to near steady-state size at 3 weeks post-infection (**Fig. 4A-B**). RNA velocity analysis (velocity length and velocity confidence; **Fig. 4C-D**) using scVelo (*10*) indicated that the velocity length of cluster 2 increased between uninfected to 1 week after *N. brasiliensis* infection, and significantly decreased from 1 week to 3 weeks after *N. brasiliensis* infection (**Fig. 4E**). The dynamics of the clusters indicate that *N. brasiliensis*-induced naïve CD4^+^ T cell heterogeneity decays over time in the absence of the helminth and is therefore a reversible event. Bulk RNAseq analysis also demonstrated that the upregulation of genes in naïve CD4^+^ T cells isolated from mice 1 week post-*N. brasiliensis* infection reversed course and was restored to near uninfected levels at 3 weeks post-infection (**Fig. 4F; Fig. S9A-B**). These transcripts included IL-4-responsive genes such as *IL4ra*, *Pros1*, *Ptpn3* and *Aqp9* (**Fig. 4F**). By contrast, type I IFN-inducible genes, which were downregulated in week 1 post-infection compared to uninfected controls, were expressed almost back at basal amounts, resembling the uninfected condition again at week 3 post-infection (**Fig. 4G**).

**Figure 4.**
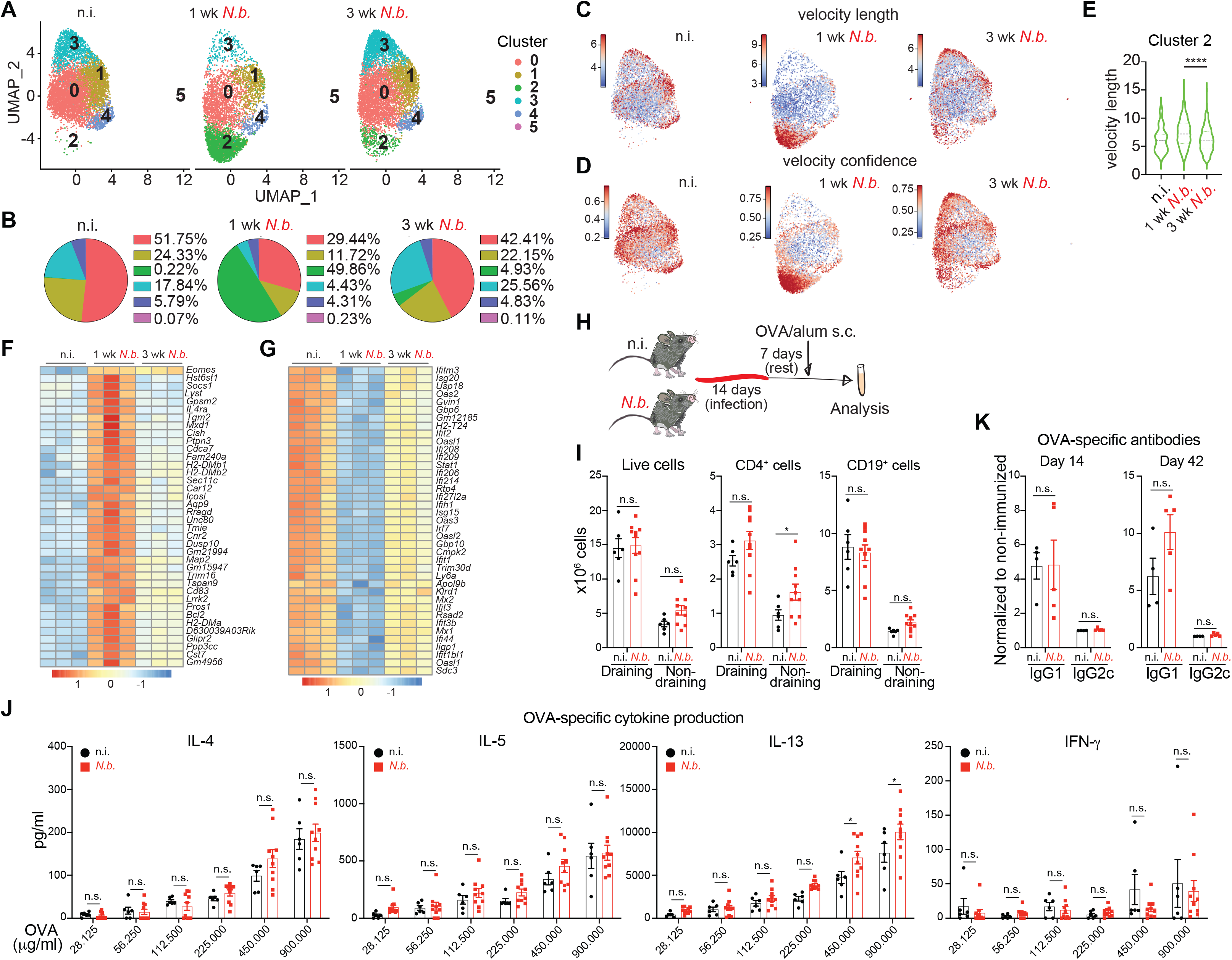
Increase in transcriptional heterogeneity and associated hypo-responsiveness of naïve T cells are reversed after helminth elimination. (**A** and **B**) scRNA sequencing of naïve CD3^+^CD4^+^CD62L^+^CD44^−^B220^−^CD25^−^CXCR5^−^PD1^−^ IL-4-GFP^−^IL-21-Kat^−^ T cells FACS-sorted from mesenteric lymph nodes (LNs) of non-infected (n.i.) mice and after 7 or 21 days post-infection with *N. brasiliensis* (1 week and 3 week *N.b*.). (**A**) Uniform Manifold Approximation and Projection (UMAP) of single naïve T cells. Each dot represents an individual cell and is colored according to cell cluster. (**B**) Pie chart depicting the percentage of naïve T cells assigned to each cluster. (**C** and **D**) Velocity length (C) and velocity confidence (D) of each cell as determined by RNA velocity. (**E**) Violin plot representing the distribution of velocity lengths of every cell assigned to cluster 2. (**F** and **G**) Heat map depicting genes upregulated (F) or downregulated (G) at 7 days, identified by RNAseq. Genes regulated at least 2-fold in naïve CD3^+^CD4^+^B220^−^CD25^−^CD44^−^CD62L^+^ T cells isolated from mesenteric LNs of n.i. mice and after 7 days post-infection with *N.b*. are shown. Gene expression is also shown for day 21 pst-infection (**H**) Experimental scheme for I to K. Mice infected with *N.b.* were immunized on day 21 post-infection with OVA/alum in the footpad. n.i. mice were also immunized as controls. (**I**) Number of total, CD4^+^CD19^−^ and CD4^−^CD19^+^ cells from draining and non-draining LNs collected 7 days post-immunization. (**J**) Concentration of indicated cytokines in the supernatant of enriched CD4^+^ T cells from draining LNs of immunized animals re-stimulated for 96 hours with OVA in the presence of irradiated splenocytes. (**K**) OVA-specific antibody concentrations for the indicated isotypes in the sera of immunized mice collected at the indicated times post-immunization. A to E are representative of 1 experiment with samples pooled from 4 mice per group where each dot represents a cell. F and G are representative of 1 experiment with three biological replicates per group. I to K are representative of 2 independent experiments and each data point represents a unique animal. Data in E were analyzed using Mann-Whitney U test. Data in I and K were analyzed using two-tailed Student’s t-test. Data in J were analyzed using 2-way ANOVA with Holm-Sidak’s multiple comparisons. ****p<0.0001 ***p<0.001, **p<0.01 and *p<0.05. n.s. is non-significant. Error bars, SEM.

To assay if the reversal of the transcriptional changes in CD4^+^ T cells correlated with a restoration of the vaccine-elicited immune response, we vaccinated uninfected mice and mice 3 weeks after *N. brasiliensis* infection (**Fig. 4H**). Mice vaccinated 3 weeks after *N. brasiliensis* infection did not exhibit changes in cellularity in either T or B cell compartments in the foot-pad draining lymph nodes compared to vaccinated, uninfected controls (**Fig. 4I**). In addition, antigen-specific CD4^+^ T cell responses, as well as OVA-specific antibody production, were also unchanged between these two groups (**Fig. 4J-K**). Unlike *N. brasiliensis* infection that is spontaneously cleared in mice by 3 weeks, *H. polygyrus bakeri* infection is chronic, lasting up to 4 months in C57BL/6 mice (*11*). Therefore, we treated *H. polygyrus bakeri*-infected mice with the anti-helminthic drug pyrantel pamoate 2 weeks post-infection. Next, we vaccinated the mice a further 2 weeks later following the treatment (**Fig. S10A**). Immunization responses elicited in these mice were indistinguishable from uninfected controls (**Fig. S10B-D**). Taken together, these results demonstrate that the IL-4-mediated transcriptional reprogramming and induced heterogeneity of naïve CD4^+^ T cell were reversible when the inducing signal *i.e.* the helminth infection was resolved. These results confirm the rate-dependent nature of naïve CD4^+^ T cell hysteresis in the context of helminth infections.

## Discussion

We discovered that naïve T cells are characterized by a cytokine-inducible, antigen-agnostic, dynamic heterogeneity that has functional consequences on their activation when subsequently encountering antigen-specific activation signals. Thus, T cell activation is subjected to pre-conditioning that can tune the magnitude of its response threshold. The activation of naïve T cells can be conceivably comparable to a state change akin to the transition between liquid to gas phases of a compound. Such a change requires the sum of exogenous activation energy similar to heat, and the average internal energy, to exceed the threshold energy required for a change in state. The membrane-bound signals for T cell activation can be considered to be analogous to the applied exogenous activation energy for phase transitions. Notwithstanding, whether an equivalence of the average internal energy state in naïve T cells exists, and if it plays a role in determining their transition to an activated state, was unknown. Our results suggest that the naïve T cell pool indeed consists of transcriptionally heterogeneous subpopulations with differential equivalence of average internal energy states for phase transition. Although intrinsic heterogeneity observed in T cells stems from lineage-specific transcriptional differences in clusters 2, 3 and 4, all lineages appear to arise independently from quiescent clusters 1 and 0. The discreteness of the type I IFN cluster *versus* the IL-4 cluster, and the inversely correlated relative changes in the size of each cluster following helminth infection, suggest a trade-off for cells to enter and commit to an individual lineage. Therefore, in the context of helminth infection and the consequent production of IL-4, the steady state equilibrium is altered to favor subpopulations with lower internal energy state equivalents. This is turn, increases the overall threshold of T cell activation. While the selective pressure for an IL-4-dependent naïve T cell threshold remains unknown, the tolerance to helminth infection, rather than helminth killing, may favor this immunosuppressive environment and higher response threshold. Specifically, IL-4 has been reported to suppress type I IFN signaling in various immune subsets including in T cells (*12*). Furthermore, administration of IL-4 prior to T cell activation suppresses antigen-specific T cell responses (*13*). The relative scaling between the type I IFN *versus* the IL-4 clusters can be expected to be bidirectional. Whether chronic infections beyond helminths, such as LCMV clone 13 infection-induced chronic T_H_1 settings, can also alter the naïve T cell threshold remains to be tested. Regardless, the steady-state heterogeneity and dynamic changes in the size of the subpopulations during various infections may pre-prime naïve T cells to shape T-dependent immune responses, independent of TCR activation.

Concurrent infections, such as overlapping helminth and other tropical diseases including malaria and tuberculosis, are known to be epidemiologically associated with dampened T_H_1 immune responses (*14–16*). Similarly, poor response to vaccination has been reported in helminth-endemic regions (*17, 18*). Helminth infection is also associated with reduced disease burden of atopic, allergic diseases and relapsing-remitting inflammatory diseases, such as inflammatory bowel disease (*19, 20*). Previously, it has been shown that factors extrinsic to antigen-specific CD4^+^ T cells, *e.g.* helminth infection-induced T regulatory cells or helminth-microbiota crosstalk-induced changes in microbiota-derived products, can impair concurrent immune responses (*21–23*). Our study reveals a novel, naïve CD4^+^ T cell-intrinsic mechanism for the integration of concurrent responses and regulation of the overall magnitude of the immune response. The reversibility of the cytokine-inducible, antigen-agnostic effect on naïve T cell heterogeneity is consistent with a rate-dependent hysteresis phenomenon where if the input is reduced to zero the output decays after a finite time lag. This is also consonant with the reversibility of helminth-mediated effects on naïve T cell heterogeneity. It has been reported that community deworming alleviated inferior immune responses observed in helminth-endemic regions of the world (*16*). However, the theoretical possibility of rate-independent responses elicited as a consequence of immune activation, perhaps brought about through permanent epigenetic changes in naïve T cells, remains. It has been demonstrated that during immune homeostasis, TGFβ epigenetically preconditions naïve CD8^+^ T cells to an epithelial resident memory phenotype (*24*). In conclusion, T cell heterogeneity is subject to inducible changes, and this dynamic heterogeneity represents a heretofore unknown mechanism for tuning immune responses.

## Acknowledgements

We are grateful to Lindsey D. Hughes for critical feedback on the manuscript. This work was supported by grants from the National Institutes of Health (NIH-NIAID R01 AI089824 and NIH-NCI R01 CA212376), Human Frontiers Science Program fellowship (LT000557/2020-L) to A.P.M., T32 (GM007205) to D.D.K. The computational analyses were carried out in part in Genifx computational facility at UAB. C.V.R is a Howard Hughes Medical Institute Faculty Scholar.

## Supplementary Materials

1. Materials and methods

2. References

3. Figs. S1 to S10

## Material and methods

### Mice

Mice were housed in specific pathogen-free conditions at Yale School of Medicine, New Haven, CT. C.129-*Il4^tm1Lky^*/J (4get) and *Il21*-*IRES-Katuska* (Kat)reporter (*Il21*^Kat/+^) mice were bred to generate *Il21*^Kat/+^*Il4*^GFP/+^ double reporter mice (*1*) and kindly provided by Dr. J. Craft (Yale University). B6.Cg-Tg(TcraTcrb)425Cbn/J (OT-II) mice were purchased from The Jackson Laboratory and subsequently bred and housed at Yale. C57BL/6J (B6) mice were purchased from The Jackson Laboratory and subsequently bred and housed at Yale. All animals were used at 8-16 weeks of age, with approval for procedures given by the Institutional Animal Care and Use Committee of Yale University.

### Helminth infections

*Nippostrongylus brasiliensis* (*N. brasiliensis*) infections were performed as previously described (2). Briefly, mice were inoculated subcutaneously with 750 viable third stage *N. brasiliensis* larvae. *Heligmosomoides polygyrus bakerii (H.p.b) larvae were a kind gift from Dr. De’Broski (University of Pennsylvania). Mice were inoculated orally with 250 viable third stage H.p.b larvae. For deworming, H.p.b-infected mice were treated on day 14 post-infection. Briefly, mice were gavaged with 7.15 mg of Strongid Paste (Zoetis) 15.25% pyrantel pamoate for a final dose of 1.09 mg of pyrantel pamoate.* Treatment efficacy was confirmed with examination of the stool for eggs 5-10 days post-treatment.

### Ovalbumin immunizations

Mice were injected subcutaneously in the footpad with 50 μL of a solution composed of 50 μg ovalbumin (OVA, Sigma-Aldrich) mixed at 1:1 ratio with Imject alum (ThermoFisher Scientific) and adsorbed by shaking for 30 minutes at room temperature. Alternatively, mice were injected subcutaneously in the footpad with 50 μL of a solution composed of 50 μg of OVA (Sigma-Aldrich) and 10 μg of ODN 1826 (CpG, Invivogen) resuspended in sterile phosphate buffered saline (PBS). Inguinal and popliteal lymph nodes (LNs) were harvested at day 7 post-immunization. Total cell numbers were counted and CD4^+^ T cells were isolated. 1×10^5^ CD4**^+^** T cells were co-cultured with 3×10^5^ irradiated splenocytes in the presence of serial dilutions of OVA beginning at 900 μg/mL. After 4 days, IL-4, IL-5, IL-13, IL-17 and IFN-γ levels were measured in the supernatant by ELISA. Additionally, at the indicated timepoints serum was collected and analyzed by ELISA for anti-OVA antibodies (Southern Biotech).

### NP-Ficoll immunization

NP-AECM-Ficoll (Biosearch Technologies) was resuspended at 1 mg/mL in sterile PBS. Mice were immunized intraperitoneally with 50 μg of NP-AECM-Ficoll 7 days post-infection with *N. brasiliensis* or in uninfected controls. Mice were bled and serum was collected 7 and 14 days post-immunization. Serum was assessed for NP-specific antibodies by ELISA.

### Labeled ovalbumin immunization

Immunizations were performed as described above but Alexa Fluor™ 488-labeled OVA (OVA-AF488, ThermoFisher Scientific) was used as the antigen. Labeled OVA was adsorbed to an equal volume of alum for 30 minutes in the dark. Mice were immunized with 50 μg of OVA in a total volume of 50 μL in the hind footpad. Additionally, one mouse per group was immunized with unlabeled OVA/alum to help establish gates for OVA-AF488 positive cells.

### Flow cytometry

Single cell suspensions were obtained in 4°C PBS with 2% fetal bovine serum (FBS) and 1 mM EDTA. Cells were incubated with anti-mouse CD16/32 Fc receptor block (clone 93, Biolegend) for 15-30 minutes. Cells were washed and stained with antibody cocktails for 30 minutes at 4°C. Live cells were labeled using Fixable Viability Dye kit (Thermofisher) according to the manufacturer’s instructions. Antibodies were purchased from commercial sources and are listed as follows: CD3 (clone 145-2C11, Biolegend), CD4 (clone RM4-5, Biolegend), CD4 (clone GK1.5, Biolegend), CD19 (clone 6D5, Biolegend), CD44 (clone IM7, Biolegend), CD45.2 (104, Biolegend), CD8a (clone 53-6.7, Biolegend), IgD (clone 11-26c.2a, Biolegend), CD138 (clone 281-2, Biolegend), B220 (clone RA3-6B2, Biolegend), PD-1 (clone 29F.1A12, Biolegend), CXCR5 (SPRL5, Thermofisher) CD86 (clone GL-1, Biolegend), CD11c (clone N418, Biolegend), MHC class II (clone M5/114.15.2, Biolegend). PE or APC-labeled streptavidin (Biolegend) was used to label biotin-conjugated antibodies again with a 30 minutes incubation at 4°C followed by washing of unbound streptavidin. Cells were washed in 1X PBS an additional time before being acquired on LSR-II (Beckton Dickinson) or Stratedigm S1000EX (Stratedigm) machines. Data were analyzed using FlowJo software (TreeStar).

### Adoptive T cell transfer and CTV labeling

Total or naïve mouse CD4^+^ T cells were isolated from the spleen or mesenteric LNs by negative selection with the EasySep Mouse CD4^+^ T cell enrichment kit or the EasySep Mouse naïve CD4^+^ T cell enrichment kit according to the manufacturer’s protocol (Stemcell Technologies). In certain cases, cells were labeled with Cell Trace™ Violet and labeled according to the manufacturer’s protocol (Thermofisher). 500,000 total or naïve T cells were then intravenously transferred to congenic CD45.1^+^ recipients.

### Enzyme-linked immunosorbent assay (ELISA)

IL-4, IL-5, IL-13, IL-17 and IFN-γ were measured by ELISA Ready-Set-Go kits from Thermofisher and culture supernatants were tested at different dilutions according to the manufacturer’s protocol. For OVA-specific ELISAs, EIA/RIA 96-well plates were coated overnight at 4°C with 200 μg/ml of OVA (Sigma-Aldrich) diluted in PBS. Serum was subsequently added and antibodies were detected using goat anti-mouse IgG1 or goat anti-mouse IgG2c conjugated to HRP (Southern Biotech) and 1xTMB (Thermofisher) or ABTS substrate (Southern Biotech).

### Fluorescence-associated cell sorting

Single cell suspensions were obtained in 4°C PBS with 2% FBS and 1 mM EDTA. Cells were incubated with anti-mouse CD16/32 Fc receptor block (clone 93, Biolegend) for 15-30 minutes. Cells were washed and stained with antibody cocktails for 30 minutes at 4°C. Antibodies were purchased from commercial sources and are listed as follows: CD3 (clone 145-2C11, Biolegend), CD4 (clone RM4-5, Biolegend), B220 (clone RA3-6B2, Biolegend), CD44 (clone IM7, Biolegend), CD62L (clone MEL-14, COMPANY), CD25 (clone PC61.5, Thermofisher), CXCR5 (clone SPRL5, Thermofisher) and PD-1 (clone J43, Thermofisher). Cells were then washed and filtered prior to sorting, which was performed using FACSARIA (BD). For the purpose of the sort, naïve CD4^+^ T cells were considered to be CD3^+^CD4^+^B220^−^IL-4-GFP^−^IL-21-Kat^−^CD25^−^CD44^−^CD62L^+^CXCR5^−^PD-1^−^ or CD3^+^CD4^+^B220^−^CD25^−^ CD44^−^CD62L^+^ and Th2 cells were considered to be CD3^+^CD4^+^B220^−^IL-4-GFP^+^IL-21-Kat^−^ CD44^+^CD62L^−^CXCR5^−^PD-1^−^.

### IL-4 blockade

Mice were injected intraperitoneally with 200 μg with anti-IL-4 (clone 11B11, Bio X Cell) or Rat IgG1 isotype control (clone HRPN, Bio X Cell). The first dose was administered 2 days prior to infection and mice were dosed every 2 days thereafter until sacrificed.

### IL-4c injection

Mice were injected with a combination of IL-4 (Peprotech) and anti-IL-4 (clone 11B11, Bio X Cell). Briefly, IL-4 was complexed with anti-IL-4 at a 1:5 mass ratio for 5 minutes at 4°C and diluted in sterile PBS. A final amount of 5 μg of IL-4 was injected intraperitoneally into mice. LN and spleen were collected after 24 hours and processed for cell sorting and RNA sequencing as described below.

### Single Cell sequencing and data processing

Reactions were performed using the Chromium Single Cell 3ʹ GEM, Library & Gel Bead Kit v3 (10x Genomics). Single cell suspensions of 20,000 FACS-sorted naïve (CD3^+^CD4^+^CD62L^+^CD44^−^B220^−^ CD25^−^CXCR5^−^PD1^−^IL-4-GFP^−^IL-21-Kat^−^) CD4^+^ T and Th2 (CD3^+^CD4^+^CD62L^−^CD44^+^B220^−^CXCR5^−^ PD1^−^IL21^−^IL4^+^) cells were added to RT Master Mix, loaded on the Single-Cell-A-Chip and partitioned with a pool of about 750,000 barcoded gel beads to form nanoliter-scale Gel Beads-In-Emulsions (GEMs). Each gel bead contained (i) an Illumina® R1 sequence (read 1 sequencing primer), (ii) a 16 nt 10x Barcode, (iii) a 12 nt Unique Molecular Identifier (UMI), and (iv) a poly-dT primer sequence (30nt). Upon dissolution of the Gel Beads in a GEM, the primers are released and mixed with cell lysate and Master Mix. Incubation of the GEMs then produces barcoded, full-length cDNA from poly-adenylated mRNA using the 10x Genomics® platform. Post GEM-RT Cleanup was performed using Silane magnetic beads to remove leftover biochemical reagents and primers from the post GEM reaction mixture. Full-length, barcoded cDNA was then amplified by PCR to generate sufficient mass for library construction. Enzymatic Fragmentation and Size Selection were used to optimize the cDNA amplicon size prior to library construction. R1 (read 1 primer sequence) were added to the molecules during GEM incubation. P5, P7, a sample index, and R2 (read 2 primer sequence) were added during library construction via End Repair, A-tailing, Adaptor Ligation, and PCR. The final libraries contained the P5 and P7 primers used in Illumina bridge amplification. Illumina-ready single Cell 3’ libraries comprised of standard Illumina paired-end constructs that begin and end with P5 and P7. The Single Cell 3’ 16 bp 10x Barcode and 12 bp UMI were encoded in Read 1, while Read 2 was used to sequence the cDNA fragment (91bp). Sequencing a Single Cell 3’ Library produced a standard Illumina BCL data output folder. The BCL data included the paired-end Read 1 (containing the 16 bp 10x™ Barcode and 12 bp UMI) and Read 2 and the sample index in the i7 index read. Minimum sequencing depth was 20,000 read pairs per cell. The output data matrix from Cell Ranger™ of individual samples were merged by *cellranger aggr* function and used to perform downstream analysis.

### Single Cell sequencing data analysis

The single cell sequencing data were analyzed by the Seurat package with the standard workflow (*3*). Briefly, cells were filtered by nFeature_RNA > 700 & nFeature_RNA < 2200 & percent.mt < 9, gene expression are normalized by “LogNormalize” and scaled for all genes. Cells were clustered with dim=1:15, resolution=0.3 and plotted on UMAP dimensions with dim=1:20. Marker genes for each cluster were calculated with FindAllMarkers function and statistical significance was calculated by Wilcoxon Rank Sum test. Single cells were randomly sampled into two individual analyses represented in Figures 1 and 4.

### Velocity Calculation

The velocity analysis was done by first generating loom files from cellranger bam output by Velocyto (*4*). The count tables of both spliced and unspliced RNA were subsetted by the cells finally present in the Seurat object. scVelo package was used for further velocity calculation (*5*). Briefly, the cluster information and UMAP embeddings were extracted from Seurat objects and added into Anadata object constructed by the count tables. The top 2000 variable genes were used for filter and normalization and summarized by scv.pp.moments(adata, n_pcs=30, n_neighbors=30). RNA velocity was calculated with a default setting of scvelo.tl.velocity function using a stochastic model and projected on the UMAP embedding. The velocity_length of cluster2 was plotted and compared using Mann-Whitney rank test.

### Cell trajectory determination

For cell-lineage determination we first resampled the data to have exactly the same number of cells in each cluster identified in Seurat. The trajectory calculation was carried out using the *Slingshot* package with Seurat clusters and default parameters (*6*). The algorithm was allowed to automatically identify the start and the end clusters. The lineages identified using *Slingshot* were evaluated visually. In addition, the diffusion map was reconstructed using the destiny package (*7*) without resampling and with 50 principle components and sigma values returned by find.sigmas and optimal.sigma functions.

### RNA Sequencing and analysis

Total RNA from sorted naïve CD4**^+^** T cells was obtained with Qiagen RNeasy Mini kit following the manufacturer’s instructions. RNA libraries were then prepared from two biological replicates per condition and samples were sequenced using 75 base pair single-end reading on HiSeq 2500 high-output instrument (Illumina). The raw reads were then subjected to trimming by Trim Galore (https://github.com/FelixKrueger/TrimGalore) with quality score 35 to remove sequencing adaptors and low-quality regions. Next, reads were mapped to the mouse genome (GRCm38) using STAR 2.7.3a with --quantMode TranscriptomeSAM (*8*) and the count table was generated by RSEM 1.3.0 rsem-calculate-expression function (*9*). Normalization and differentially expressed genes (DEGs) were calculated by DEseq2 package (*10*). DEGs with an adjusted p-value lower than 0.05 were used generating volcano plots with EnhancedVolcano package (cutoff line padj<10^10^ and |Log2FoldChange|>1). For the IL-4 neutralization experiment, the normalization was done using EdgeR package (*11*). DEGs with higher than two-fold change (both up- and down-regulated) were used in making a Venn plot using the VennDiagram package. The hypergeometric p value was calculated using the gmp package with a total gene number of 44791. Heatmaps were plotted using pheatmap package and expression values were scaled by row to visualize the lowly expressed genes.

### Nanostring Analysis

Adoptively transferred CD45.2^+^CD4^+^ OT-II T cells were FACS-sorted from draining lymph nodes 4 days post immunization and lysed in RLT lysis buffer diluted in water. Cell lysates were processed according to the manufacturer’s recommendations using the nCounter PanCancer Immune Profiling Panel and the nCounter system (NanoString Technologies). The data were subsequently analyzed using version 4.0 of the nSolver™ Analysis Software (NanoString Technologies) where the samples were normalized based on the panel of housekeeping genes as recommended by the manufacturer using the geoNorm algorithm (*12*). Only genes that were significantly different (p < 0.05; t test, unadjusted) between groups were considered. Pathway scores were generated using the first principal component of the gene expression data as established by Tomfohr *et al. (13)*.

### RT-quantitative PCR

Reverse transcription of RNA was performed utilizing iScript cDNA Synthesis Kit (Bio-Rad Laboratories) according to the manufacturer’s instructions. cDNA fragments were then amplified using KAPA SYBR Fast qPCR Kit (Kapa Biosystems) according to the manufacturer’s instructions and qPCR reactions were carried out on CFX96 Thermal Cycler Real Time System (Bio-Rad Laboratories). The reactions were normalized to housekeeping genes (m*Rlp13a* and m*S18*) and the specificity of the amplified products was verified by dissociation curves. The following primers were purchased for use from Sigma: m*Aqp9* fwd 5′-TGGTGTCTACCATGTTCCTCC-3′, m*Aqp9* rev 5′- AACCAGAGTTGAGTCCGAGAG-3′; m*Lgals3* fwd 5′-AGACAGCTTTTCGCTTAACGA-3′, m*Lagls3* rev 5′-GGGTAGGCACTAGGAGGAGC-3′; m*Ppp3cc* fwd 5′-ATGCCACCCCGAAAAGAGG-3′, m*Ppp3cc* rev 5′-CATGGTCGGTCCTTCTTGACG-3′; m*Pros1* fwd 5′-GCACAGTGCCCTTTGCCT-3′, m*Pros1* rev 5′-CAAATACCACAATATCCTGAGACGTT-3′; m*Rlp13a* fwd 5′- CTGTGAAGGCATCAACATTTCTG-3′, m*Rlp13a* rev 5′-GACCACCATCCGCTTTTTCTT-3′; m*S18* fwd 5′-AGTTCCAGCACATTTTGCGAG-3′, m*S18* rev 5’-TCATCCTCCGTGAGTTCTCCA-3′; m*Tgm2* fwd 5′-GACAATGTGGAGGAGGGATCT-3′, m*Tgm2* rev 5′-CTCTAGGCTGAGACGGTACAG-3′.

### Cluster Similarity Analysis

The similarity between the naïve T cell clusters in our work and those of ElTanbouly *et al.* (*14*) was calculated using the Jaccard similarity among the clusters. This was calculated as a ratio of intersection and union of the marker gene symbols as identified by Seurat. We calculated the pairwise Jaccard similarity of all clusters identified by two studies. We determined the P-value of this similarity measure using a permutation test where all cluster labels were shuffled 1000 times and Jaccard similarities derived from this shuffled data were used to assign p-values to the observed similarity. The final identification of clusters was done using the maximum similarity value that passed the p-value cutoff of 0.05.

### Statistical analysis

All statistical analyses were done using GraphPad Prism (GraphPad Software Inc.). Data are shown as mean ± S.E.M. and each data point represents a unique animal or pooled samples as indicated in respective figure legends. Statistical differences between two experimental groups were determined by two-tailed Student’s t-test. Statistical differences between three or more experimental groups were determined by two-way ANOVA followed by Holm-Sidak’s multiple comparisons test. Significance was defined as *p<0.05, **p<0.01, ***p<0.001, and ****p<0.0001. n.s. is non-significant.

---

**Figure S1.**
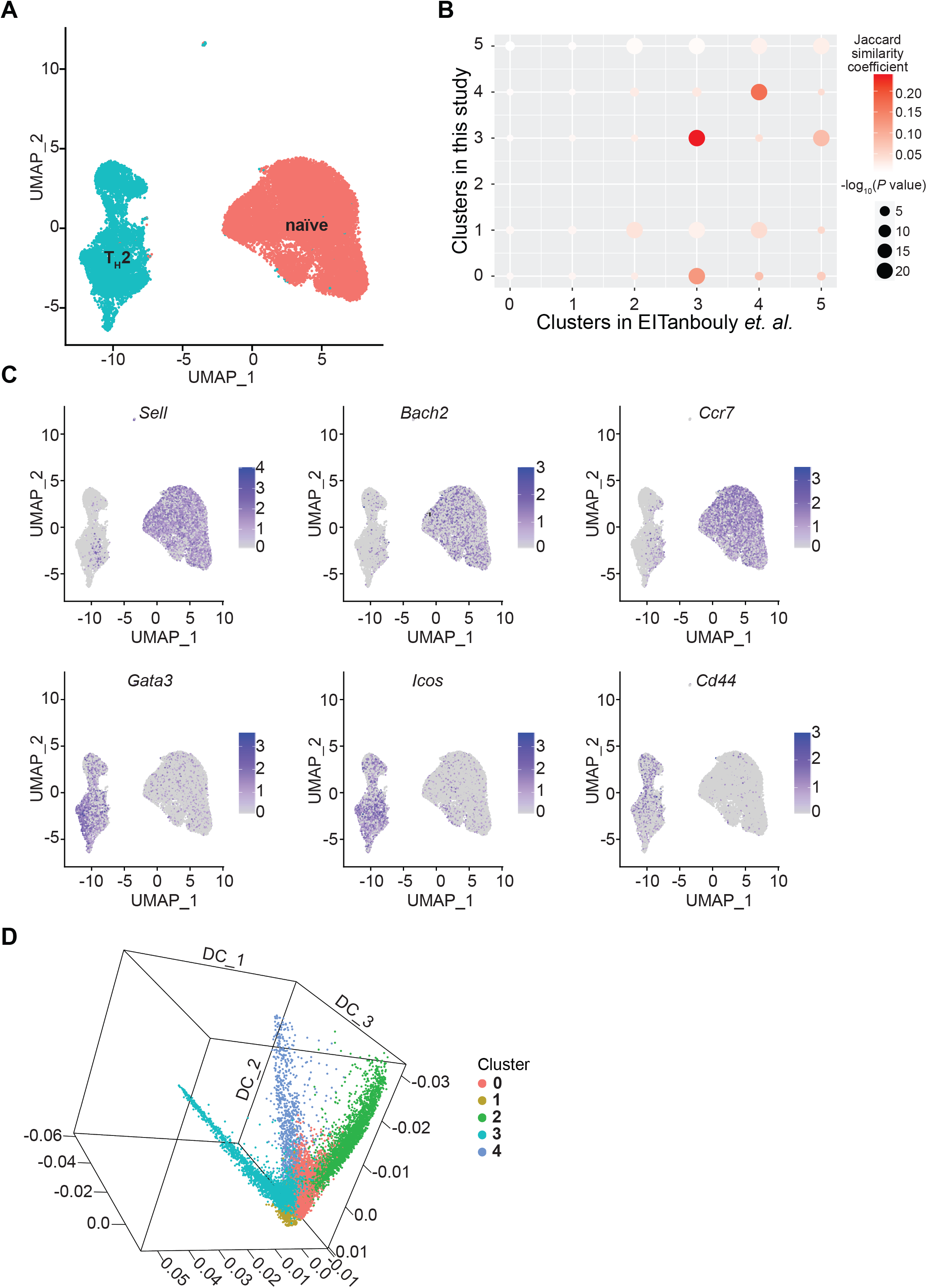
Naïve T cell clusters are distinct from Th2 cells during helminth infection. scRNA-seq was performed on CD3^+^CD4^+^CD62L^+^CD44^−^B220^−^CD25^−^CXCR5^−^PD1^−^IL-4-GFP^−^IL-21-Kat^−^ naïve T cells and CD3^+^CD4^+^CD62L^−^CD44^+^B220^−^CXCR5^−^PD1^−^IL-4-GFP^+^IL-21-Kat^−^ Th2 cells FACS-sorted from mesenteric lymph nodes (LNs) of *N.b.*-infected (day 7) mice. (**A**) Uniform Manifold Approximation and Projection (UMAP) of single naïve T and Th2 cells. Each dot corresponds to one cell where red dots represent sorted naïve T cells and cyan dots represent sorted Th2 cells. (**B**) Cluster similarity was performed by calculating the pairwise Jaccard similarity of all clusters identified in the naïve T cell dataset included in Figure 1 and that of ElTanbouly *et al.* (**C**) Expression of indicated genes by single naïve T and Th2 cell clusters depicted by UMAP. Each dot corresponds to one cell. (**D**) 3D scatter plot showing the 3 trajectories identified by diffusion map. x, y, and z axes represent diffusion components 1, 2 and 3, respectively. Dots are cells and colored as in Figure 1C. Cluster 2, 3 and 4 were found in distinct trajectories. A to D are representative of one experiment with samples pooled from 4 mice per group.

**Figure S2.**
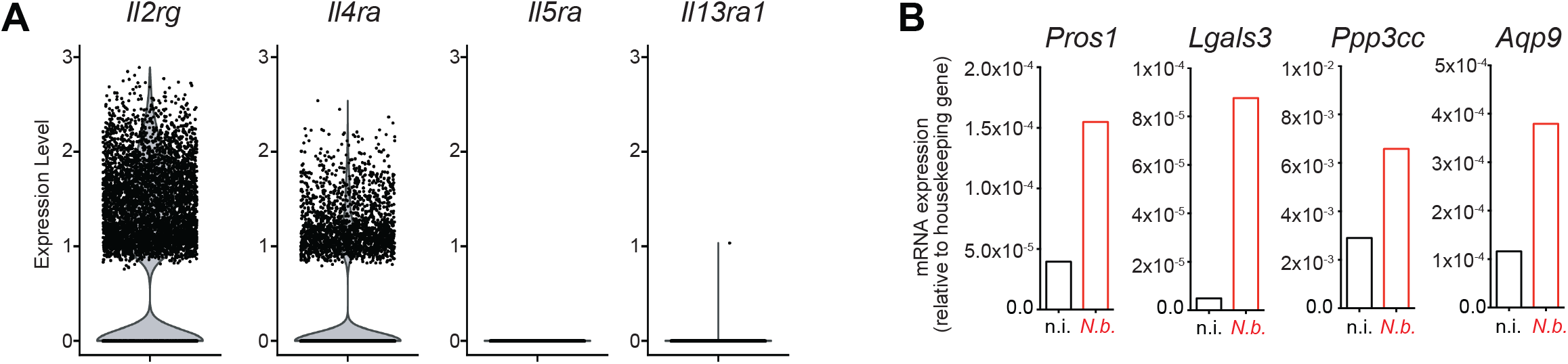
Naïve CD4^+^ T cells express the IL-4 receptor subunits and display transcriptional changes upon *N.b.* infection. (**A**) Violin plots representing the distribution of normalized expression levels of indicated cytokine receptor genes from scRNA-seq performed on FACS-sorted CD3^+^CD4^+^CD62L^+^CD44^−^B220^−^CD25^−^ CXCR5^−^PD1^−^IL-4-GFP^−^IL-21-Kat^−^ naïve T cells from mesenteric lymph nodes (LNs) of *N.b.*-infected (day 7) mice. Representative of 1 experiment with samples pooled from 4 mice. (**B**) mRNA expression levels for indicated genes measured by RT-qPCR in FACS-sorted naïve CD3^+^CD4^+^CD62L^+^CD44^−^B220^−^CD25^−^CXCR5^−^ PD1^−^IL-4-GFP^−^IL-21-Kat^−^ T cells from mesenteric LNs of non-infected (n.i.) and *N.b.*-infected (day 7) mice. Representative of 2 independent experiments with samples pooled from 4 mice per group.

**Figure S3.**
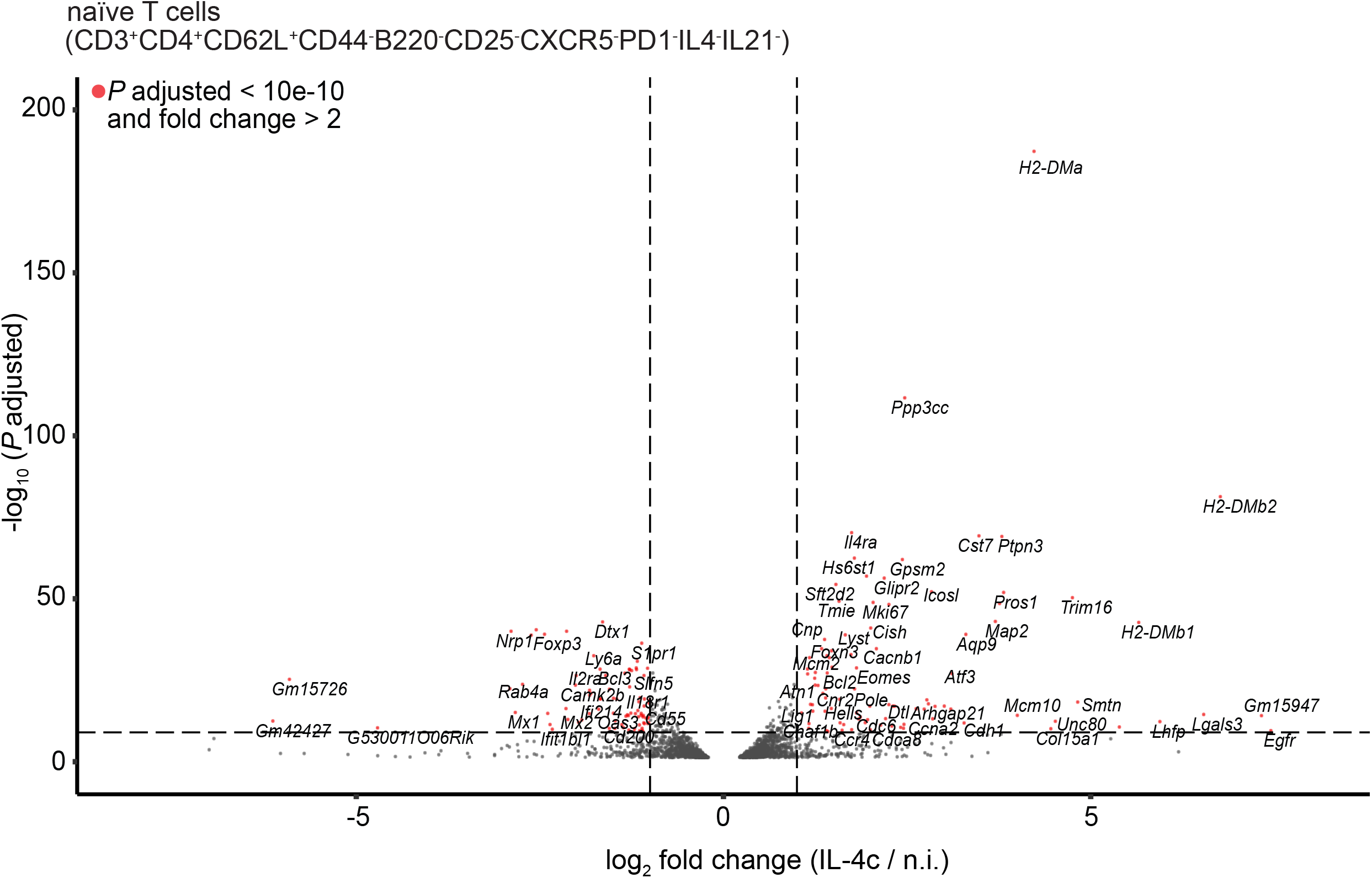
IL4c treatment results in transcriptional changes in naïve CD4^+^ T cells. Naïve CD3^+^CD4^+^CD62L^+^CD44^−^B220^−^CD25^−^CXCR5^−^PD1^−^IL-4-GFP^−^IL-21-Kat^−^ T cells were FACS-sorted from mesenteric lymph nodes from IL-4c-treated mice 1 day post-injection or untreated controls. Volcano plot represents genes differentially regulated at least 2-fold in IL-4c-treated mice in comparison to controls, as determined by RNAseq. Representative of 2 independent experiments with samples pooled from 4 mice per group.

**Figure S4.**
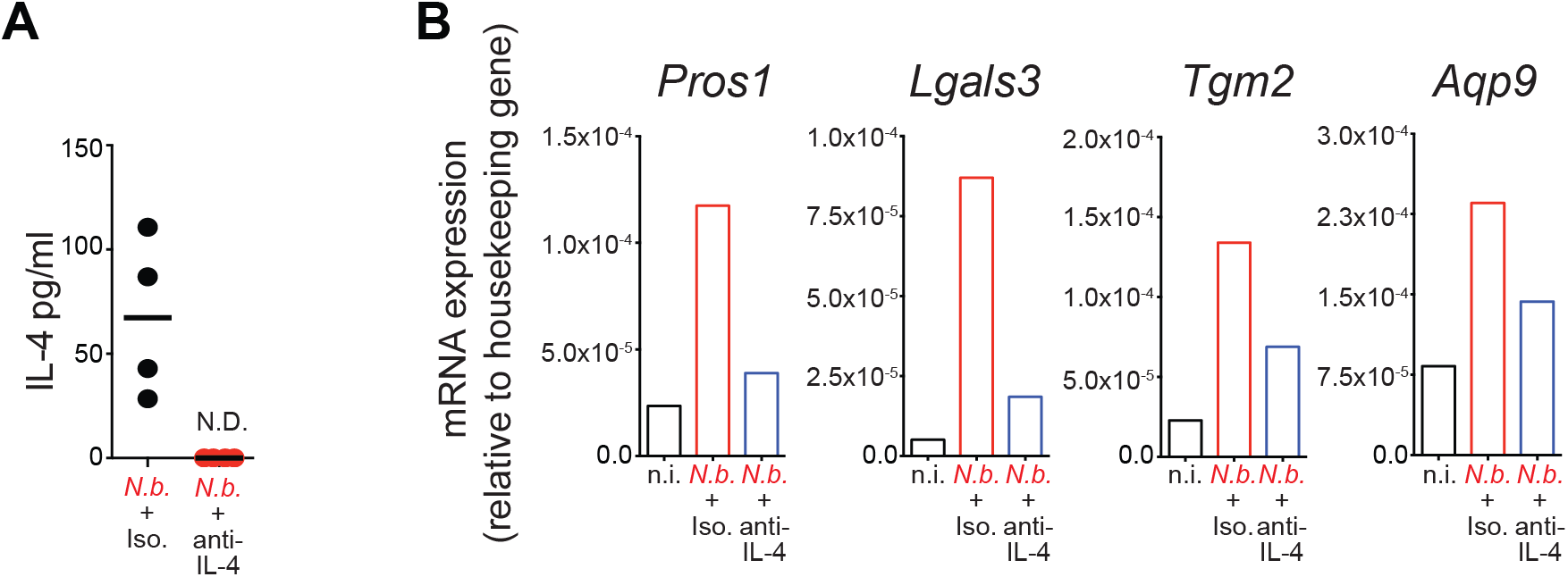
IL-4 blockade suppresses helminth infection-induced genes in naïve T cells. *N.b.*-infected mice were treated with 200μg IL-4 neutralizing antibody or isotype control beginning 48 hours prior to *N.b.* infection and every 48 hours thereafter until mice were sacrificed at 7 days post-infection. (**A**) Serum concentration of IL-4 in mice administered isotype or IL-4 neutralizing antibody as determined by ELISA using the same antibody clone as the IL-4 neutralizing antibody. Each data point represents a unique animal. (**B**) Expression of indicated genes in FACS-sorted naïve CD3^+^CD4^+^CD62L^+^CD44^−^B220^−^CD25^−^ T cells from mesenteric lymph nodes of non-infected (n.i.), *N.b.*-infected plus isotype-treated (day 7) and *N.b.*-infected plus anti-IL-4 antibody-treated (day 7) mice, as determined by RT-qPCR. Representative of 1 experiment with samples pooled from 4 mice per group.

**Figure S5.**
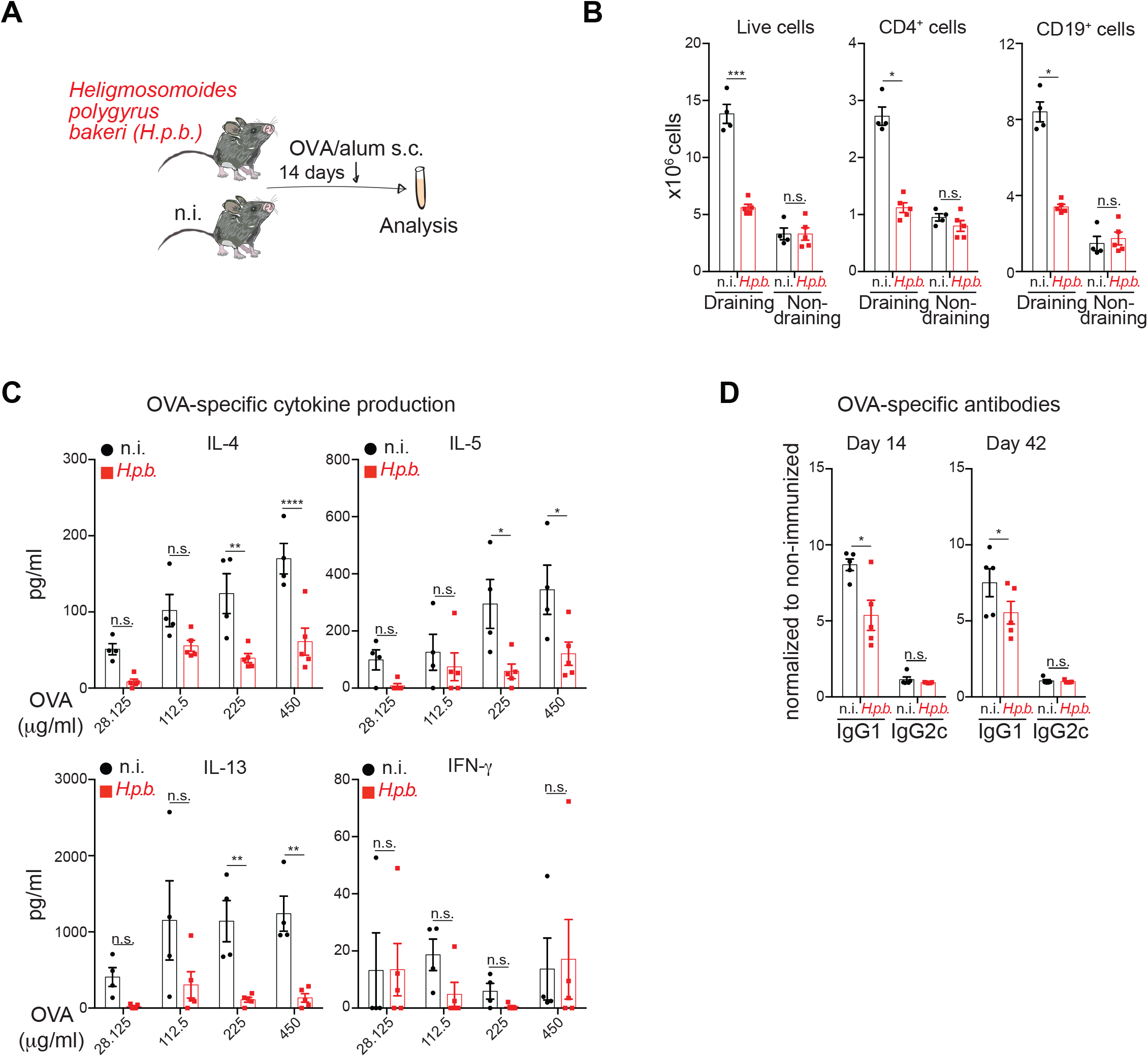
*H.p.b.* infection impairs antigen specific responses after OVA-alum immunization. (**A**) Experimental scheme for B to D. Non-infected (n.i.) controls or mice infected with *Heligmosomoides polygyrus bakeri* (*H.p.b.*) for 14 days were immunized with OVA/alum subcutaneously in the footpad. (**B**) Number of total, CD4^+^CD19^−^ and CD4^−^CD19^+^ cells from draining and non-draining lymph nodes (LNs) collected 7 days post-immunization. (**C**) Concentrations of indicated cytokines in the supernatant of enriched CD4^+^ T cells from draining LNs of immunized animals re-stimulated for 96 hours with OVA in the presence of irradiated splenocytes. (**D**) OVA-specific antibody concentrations for the indicated isotypes in the sera of immunized mice collected at the indicated times post-immunization. Representative of 3 independent experiments and each data point represents a unique animal. Data in B and D were analyzed using two-tailed Student’s t-test. Data in C were analyzed using 2-way ANOVA with Holm-Sidak’s multiple comparisons test. ****p<0.0001 ***p<0.001, **p<0.01 and *p<0.05. n.s. is non-significant. Error bars, SEM.

**Figure S6.**
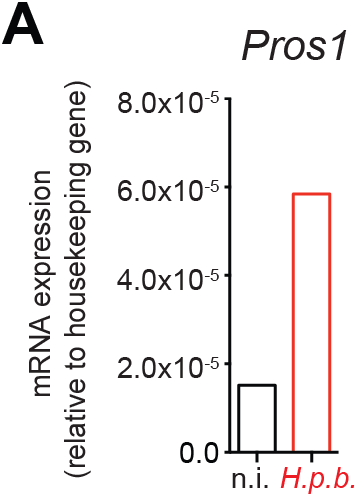
*H.p.b*. infection results in the upregulation of an IL-4 inducible gene in naïve T cells. Naïve CD3^+^CD4^+^CD62L^+^CD44^−^B220^−^CD25^−^ T cells were FACS-sorted from mesenteric lymph nodes of *H.p.b.*- infected (day 14) or non-infected (n.i.) mice. *Pros1* expression was determined by RT-qPCR. Representative of 2 independent experiments with samples pooled from 4 mice per group.

**Figure S7.**
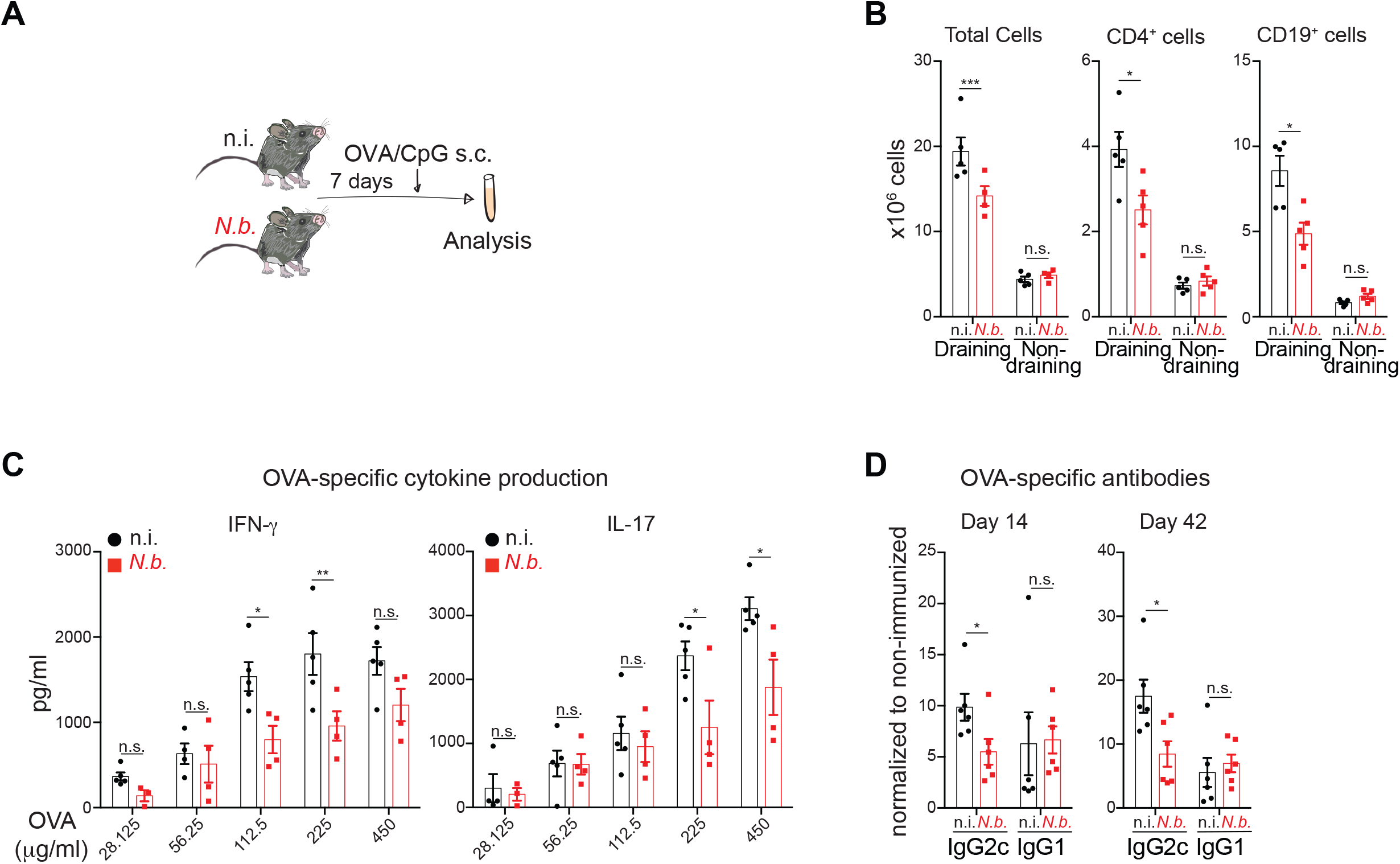
*N.b.* infection suppresses antigen-specific type 1 and 17 immune responses to OVA/CpG immunization. (**A**) Experimental scheme for B to D. Non-infected (n.i.) controls or mice infected with *N.b.* for 7 days were immunized with OVA and CpG in the footpad. (**B**) Number of total, CD4^+^CD19^−^ and CD4^−^CD19^+^ cells from draining and non-draining lymph nodes (LNs) collected 7 days post-immunization. (**C**) Concentrations of indicated cytokines in the supernatant of enriched CD4^+^ T cells from draining LNs of immunized animals re-stimulated for 96 hours with OVA in the presence of irradiated splenocytes. (**D**) OVA-specific antibody concentrations for the indicated isotypes in the sera of immunized mice collected at the indicated times post-immunization. Representative of 2 independent experiments and each data point represents a unique animal. Data in B and D were analyzed using two-tailed Student’s t-test. Data in C were analyzed using 2-way ANOVA with Holm-Sidak’s multiple comparisons test. ****p<0.0001 ***p<0.001, **p<0.01 and *p<0.05. n.s. is non-significant. Error bars, SEM.

**Figure S8.**
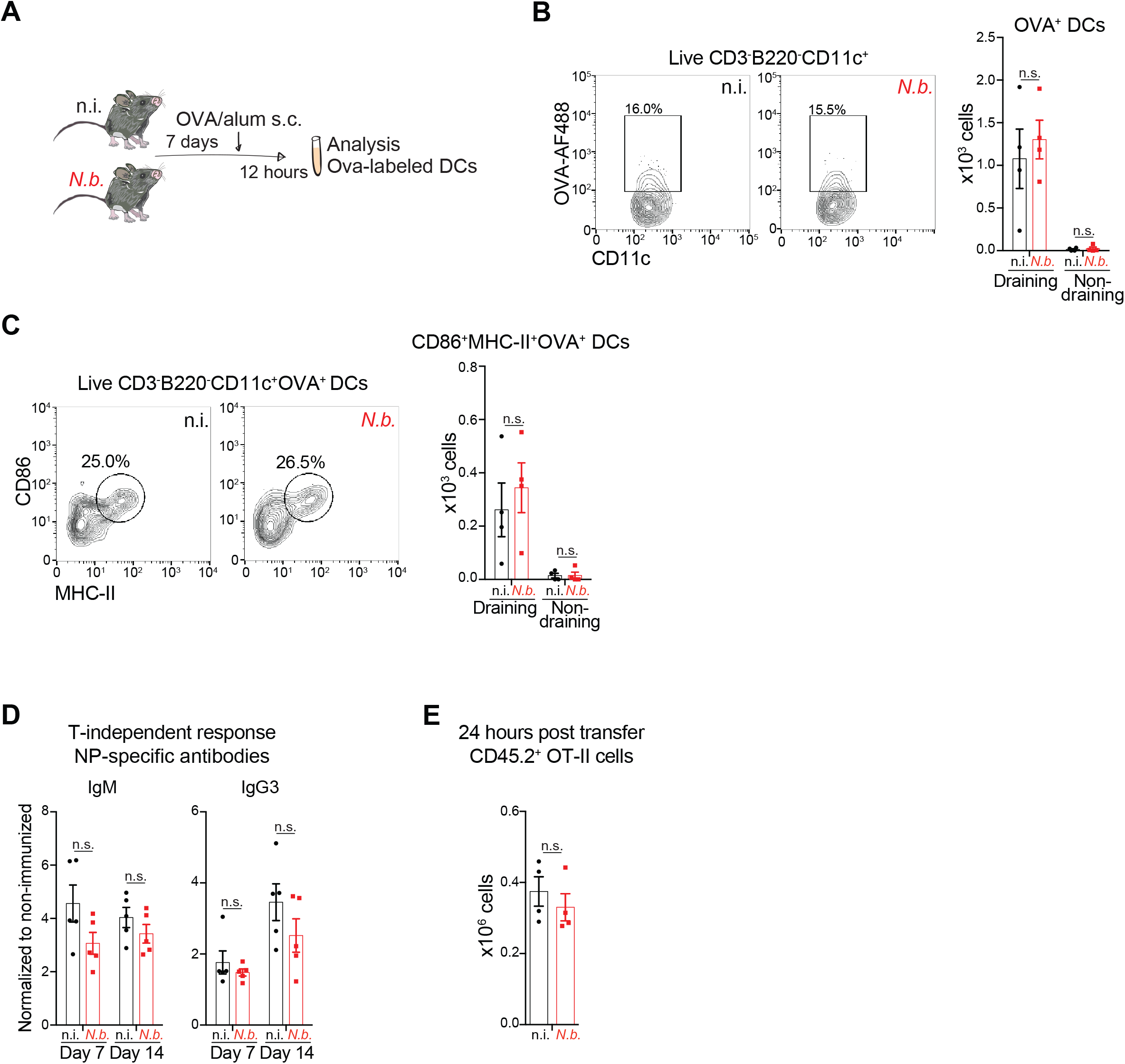
*N.b.* infection does not impact DC antigen delivery, T-independent B cell responses and naïve T cell circulation. (**A**) Experimental scheme for B and C. Non-infected (n.i.) controls or mice infected with *N.b.* for 7 days were injected with fluorescently-labeled ovalbumin(OVA-AF488)/alum in the footpad. Draining and non-draining lymph nodes (LNs) were collected 12 h post-immunization. (**B**) Representative contour plot and total cell counts of OVA^+^CD3^−^B220^−^CD11c^+^ dendritic cells (DCs) from animals immunized as in A. (**C**) Representative contour plot and total cell counts of OVA^+^CD86^+^MHCII^+^CD3^−^B220^−^CD11c^+^ DCs. (**D**) N.i. controls or mice infected with *N.b.* for 7 days were immunized intra-peritoneally with NP-Ficoll. Serum was collected at the indicated times post-immunization and NP-specific antibody levels were measured. (**E**) N.i. CD45.1^+^ control or CD45.1^+^ mice infected with *N.b.* for 7 days received 500,000 CD45.2^+^CD4^+^ OT-II T cells. Numbers of CD45.2^+^CD4^+^ OT-II T cells in popliteal and inguinal LNs 24 h post-transfer into respective congenic CD45.1 recipient mice are represented. Representative of 2 independent repeats and each data point represents a unique animal. Data in B to E were analyzed using two-tailed Student’s t-test. ****p<0.0001 ***p<0.001, **p<0.01 and *p<0.05. n.s. is non-significant. Error bars, SEM.

**Figure S9.**
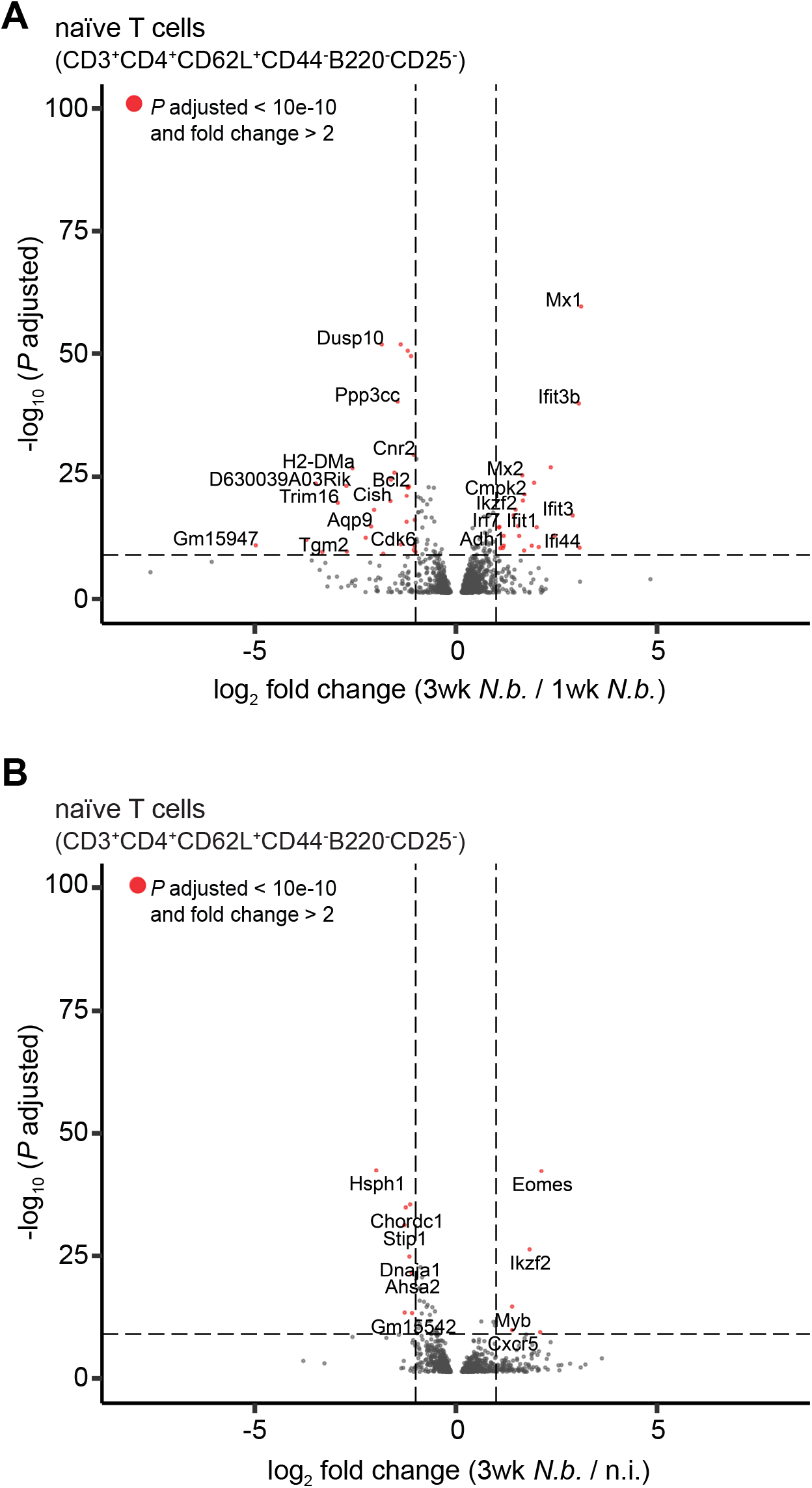
*N.b.* infection results in rate-dependent transcriptional changes in naïve CD4^+^ T cells. Naïve CD3^+^CD4^+^CD62L^+^CD44^−^B220^−^CD25^−^ T cells were FACS-sorted from mesenteric lymph nodes of *N.b.*-infected (day 7), *N.b.*-infected (day 21) or non-infected (n.i.) mice. Volcano plot representing genes differentially regulated at least 2-fold in (**A**) 3 wk *N.b.-*infected mice compared to 1 wk *N.b.*-infected mice and (**B**) 3 wk *N.b.*- infected mice compared to n.i. mice. Representative of 1 experiment with three biological replicates per group.

**Figure S10.**
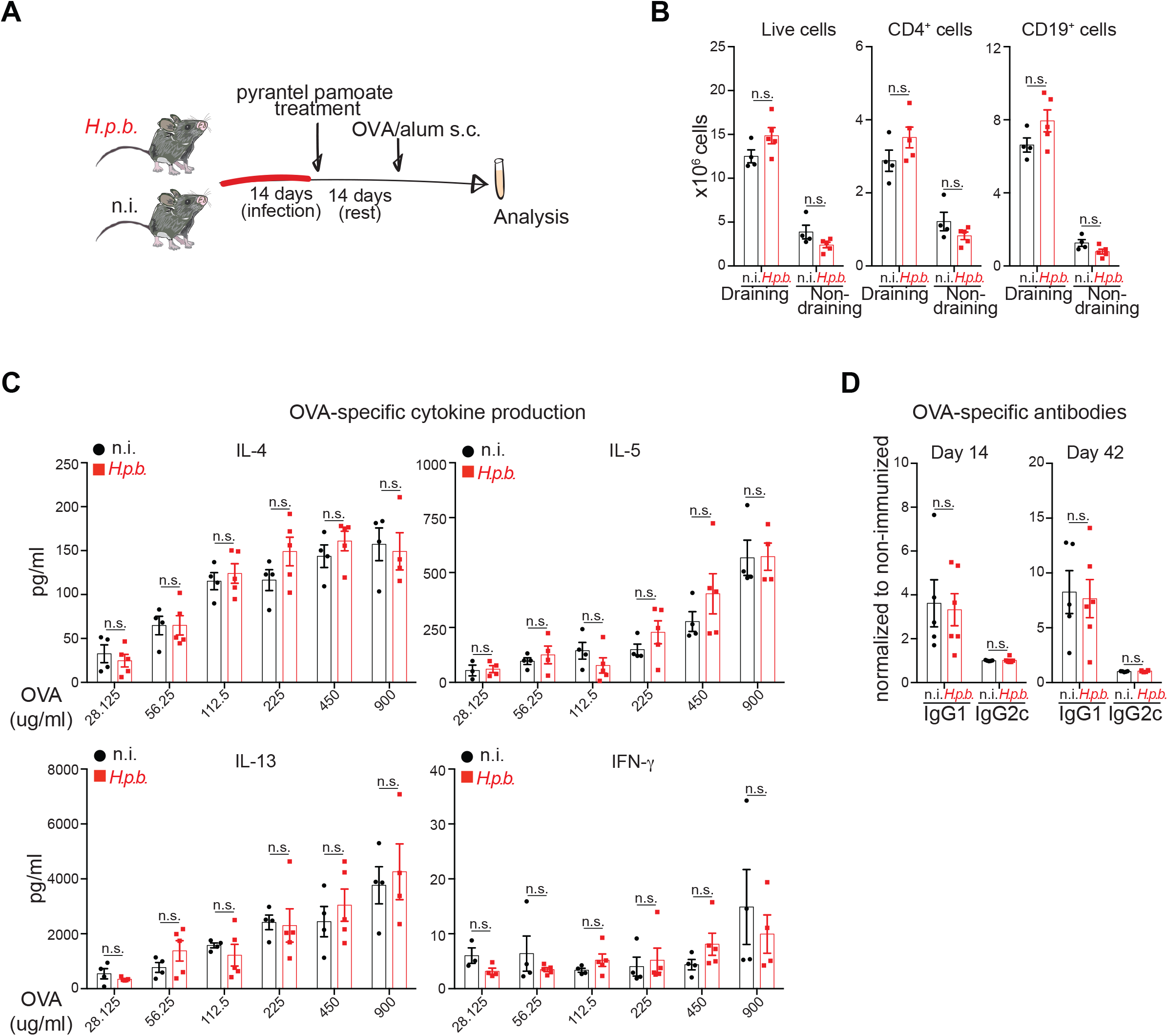
*H.p.b.* clearance correlates with restored antigen-specific immunization responses. (**A**) Experimental scheme for B to E. Mice infected with *H.p.b.* or non-infected (n.i.) controls were treated with pyrantel pamoate at 14 days post-infection and subsequently immunized 14 days post-treatment with OVA/alum in the footpad. (**B**) Number of total, CD4^+^CD19^−^ and CD4^−^CD19^+^ cells from draining and non-draining lymoh nodes (LNs) collected 7 days post-immunization. (**C**) Concentration of indicated cytokines in the supernatant of enriched CD4^+^ T cells from draining LNs of immunized animals re-stimulated for 96 hours with OVA in the presence of irradiated splenocytes. (**D**) OVA-specific antibody concentrations for the indicated isotypes in the sera of immunized mice collected at the indicated times post-immunization. Representative of 2 independent experiments and each data point represents a unique animal. Data in B and D were analyzed using two-tailed Student’s t-test. Data in C were analyzed using 2-way ANOVA with Holm-Sidak’s multiple comparisons test. ****p<0.0001 ***p<0.001, **p<0.01 and *p<0.05. n.s. is non-significant. Error bars, SEM.

## Notes

### Competing Interest Statement

The authors have declared no competing interest.

